# Spatial distribution of local patch extinctions drives recovery dynamics in metacommunities

**DOI:** 10.1101/2020.12.03.409524

**Authors:** Camille Saade, Sonia Kéfi, Claire Gougat-Barbera, Benjamin Rosenbaum, Emanuel A. Fronhofer

## Abstract

Human activities lead more and more to the disturbance of plant and animal communities with local extinctions as a consequence. While these negative effects are clearly visible at a local scale, it is less clear how such local patch extinctions affect regional processes, such as metacommunity dynamics and the distribution of diversity in space. Since local extinctions may not be isolated events in space but rather cluster together, it is crucial to investigate their effects in a spatially explicit framework.

Here, we use experimental microcosms and numerical simulations to understand the relationship between local patch extinctions and metacommunity dynamics. More specifically, we investigate the effects of the amount and spatial autocorrelation of extinctions in a full factorial design. Experimentally, we found that local patch extinctions increased inter-patch (*β*-) diversity by creating differences between perturbed and unperturbed patches and at the same time increased local (*α*-) diversity by delaying the competitive exclusion of inferior competitors. Most importantly, recolonization dynamics depended more strongly on the spatial distribution of patch extinctions than on the amount of extinctions per se. Clustered local patch extinctions reduced mixing between perturbed and unperturbed patches which led to slower recovery, lower *α*-diversity in unperturbed patches and higher *β*-diversity. Results from a metacommunity model matched the experimental observations qualitatively when the model included ranked competitive interactions, giving a hint at the underlying mechanisms.

Our results highlight that local patch extinctions can increase the diversity within and between communities, that the strength of these effects depends on the spatial distribution of extinctions and that the effects of local patch extinctions can spread regionally, throughout a landscape. These findings are highly relevant for conservation and management of spatially structured communities under global change.

## Introduction

Understanding the causes and consequences of local extinctions and how they affect biological systems at larger spatial scales lies at the heart of spatial ecology. Natural metapopulations and metacommunities — sets of local populations and communities linked by dispersal (Levins, 1969) — naturally experience local extinctions (Altermatt and Ebert, 2010; Fronhofer, Kubisch, et al., 2012; Hanski and Kuussaari, 1995), for instance, due to demographic stochasticity, natural disasters or disease outbreaks. In addition, global changes — including climate change, habitat loss and fragmentation due to land-use changes, deforestation and urbanization — put increasing stress on ecological communities (IPBES, 2019; Millennium Ecosystem Assessment, 2005) which contributes to local patch extinctions.

Local patch extinctions, which we here define as the disappearance of all species from a patch, can have various consequences. In trophic systems, sustained local patch extinctions can induce regional species extinctions (Liao et al., 2017; Ryser et al., 2019) and thus reduce regional diversity. Top predators are more likely to go extinct than intermediate and basal species. As a consequence, prey species can even benefit at the regional scale from local patch extinctions due to the release from predation pressure. Furthermore, microcosms experiments on a competitive community with a competition-colonization trade-off show that occasional local patch extinctions can prevent regional extinctions and increase regional diversity by allowing less competitive species to persist (Cadotte, 2007).

One important factor mitigating the effect of local patch extinctions is the fact that metacommunities consist of independent units, the patches harbouring local communities, that are linked in space by dispersal events. The coupling of spatially distinct communities can reduce the effect of local extinctions if individual local communities face them at different times: patches left empty by a local extinction event can be recolonized through dispersal of individuals from patches that are occupied. However, dispersal between local communities can also have detrimental effects by synchronizing populations and thereby decreasing spatial insurance effects (Abbott, 2011). Under strong dispersal, the effects of local extinctions can even spread throughout a metacommunity such that local events have a regional effect (Gilarranz et al., 2017; Zelnik et al., 2019).

One likely important factor that modulates the effects discussed above is the spatial distribution of local patch extinctions, for instance, whether they are clustered in space or not. An increase in the spatial autocorrelation of local extinction events could have a destabilizing effect at the metacommunity scale by coupling local dynamics and thus increasing global extinction risk (Kahilainen et al., 2018; Ruokolainen, 2013). Indeed, climate models have predicted an increase in the spatial and temporal autocorrelation of temperature (Di Cecco and Gouhier, 2018), implying an increase in the environmental similarity between communities in space and time. This is expected to result in more climate extremes, such as heatwaves, droughts or frosts, affecting increasingly larger areas and for a longer time. Such climatic extremes can lead to local extinctions of populations of organisms sensitive to temperature changes, as seen in episodes of coral bleaching (Carpenter et al., 2008) or forest die-offs (Allen et al., 2010).

Despite this trend of climate data and predictions showing an increase in spatial and temporal correlation of temperature (Di Cecco and Gouhier, 2018) that could result in a greater number of climate-induced local extinctions and a stronger spatial autocorrelation of these events, few studies have considered the spatial structure and extent of local extinctions, leaving a gap in our understanding of how spatially clustered extinctions may affect the dynamics of ecological systems.

Here, we investigate how the amount and spatial distribution of local patch extinctions affect recolonization dynamics in metacommunities. We were particularly interested in determining whether the effects of local patch extinctions can spread in space and have regional effects on metacommunities. Using a full factorial design crossing three levels of extinction amounts and two levels of spatial autocorrelation, we forced local patch extinctions in experimental and simulated metacommunities and followed community dynamics in each patch. We focused on the dynamics of the recolonization process (*i.e.*, during the two weeks following the extinctions) to capture the transient effects of extinctions. We were able to show that the effects of local patch extinctions on the metacommunity depend more on the spatial distribution of those extinctions than on their amount, and that local patch extinctions can increase both local (*α*-) and inter-patch (*β*-) diversity.

## Material and methods

We used a combination of laboratory experiments with metacommunities of three freshwater ciliates (*Tetrahymena thermophila*, *Colpidium* sp. and *Blepharisma* sp.) in microcosm landscapes and mathematical modelling of metacommunities to address our main research question. To do so, we forced local patch extinctions (not sustained in time, i.e., ‘pulse’ perturbations; see Bender et al. 1984) in experimental microcosm landscapes (Altermatt, Fronhofer, et al., 2015) and followed metacommunity recovery in terms of species diversity and biomass as a function of the intensity (amount of extinctions) and spatial distribution (clustered vs. dispersed) of the extinctions. Experiments and simulations followed the dynamics of metacommunities in landscapes made of 16 patches arranged in a square lattice and connected by active dispersal.

### Experiments

We used experimental landscapes made of 16 vials connected to their 4 nearest neighbours, allowing individuals to disperse from one patch to another. Local patch extinctions consisted in removing all individuals of all species in a given patch. Each patch was initially inoculated with one of the three species at half its carrying capacity. Extinctions were implemented once, two weeks after inoculation to allow for community assembly to have taken place. Subsequently, we observed the recovery of the landscapes. Since we expected the extinctions to have only a transient effect before the metacommunity reached an equilibrium dominated by the best competitor (*Blepharisma* sp.), we followed the recovery dynamics just after the extinctions for a duration of two weeks (which is the time it takes for *Blepharisma* sp. to exclude the other species in a single patch co-culture; Fig. S5 h-j). In order to explore the effects of the amount of local patch extinctions and their spatial autocorrelation on the dynamics of metacommunities, we used a full factorial design crossing three levels of local patch extinctions (0, 4 or 8 simultaneous extinctions out of 16 patches) with two levels of spatial autocorrelation (clustered: Fig. S1 landscapes 7-9 and 13-15; dispersed: Fig. S1 landscapes 4-6 and 10-12).

This design yielded a total of 5 treatments (no extinction, 4 clustered extinctions, 4 dispersed extinctions, 8 clustered extinctions, 8 dispersed extinctions) that were each replicated in 3 landscapes, for a total of 15 landscapes and 240 patches. We followed the metacommunity dynamics through time by measuring the density of each species in each patch three times per week using video recording and analysis.

#### Species

We used three freshwater ciliate species commonly used in microcosms experiments (Cadotte, 2006; Diehl and Feissel, 2001; Worsfold et al., 2009): *Tetrahymena thermophila* (Tet) is a small (50 μm, Fig. S2) bacterivore, *Colpidium* sp. (Col) is a medium-sized (120 μm, Fig. S2) bacterivore and *Blepharisma* sp. (Ble) is a big (200 μm, Fig. S2) omnivore feeding on bacteria and smaller ciliates. In this experimental system, all three species feed on the bacterium *Serratia marcescens* as a common resource and thus constitute a competition network. In addition, the biggest *Blepharisma* sp. individuals may also feed on *T. thermophila*. We determined the species’ demographic traits in preliminary single patch experiments: the species show differences in population growth rate (Tet > Col > Ble), carrying capacity (Tet > Col > Ble; Fig. S5 a-c) and interspecific competitive ability (Tet < Col < Ble; Fig. S5 h-j). Based on their population growth rates and competitive abilities, these species can be described as an ecological succession: *T. thermophila* density peaks after approximately two days, *Colpidium* sp. density peaks after approx. five days and *Bleparisma* sp. grows slowly and dominates the community after around 16 days (Fig. S5 h-j) in our experimental setting.

We did not quantify dispersal in isolation, but used movement speed observed *in situ* as a proxy of dispersal ability, as these two traits are usually well correlated (Fronhofer and Altermatt, 2015; Pennekamp, Clobert, et al., 2019). Generally, *Colpidium* sp. is faster than both *T. thermophila* and *Blepharisma* sp., which move at roughly the same speed (Fig. S3).

#### Culture conditions

The species were kept in 20 mL of a standardized medium made of water (Volvic), dehydrated organic salad (1 g of salad for 1.6 L of water) and bacteria (*Serratia marcescens*) at 10% of their maximum density (obtained by a tenfold dilution of a one week old culture) as a common resource. The cultures were refreshed three times per week by replacing 2 mL of each culture with 2 mL of fresh, bacterized medium. The cultures were kept in a room with controlled temperature (20 °C). In order to exclude any potential confounding effects due to landscape positioning, the position and orientation of landscapes was randomized and changed three times per week.

#### Landscape design

We used landscapes made of 16 vials (20 mL Sarstedt tubes) arranged in a square lattice and connected by silicon tubes (length: 6 cm, inner diameter: 4 mm). The silicon tubes were closed with clamps to control dispersal. The clamps were opened for 4 hours three times per week (after medium replacement) to allow dispersal. Each patch was initially inoculated with one of the three species at half of its carrying capacity at the beginning of the experiment. Initial species distributions were drawn at random so that one species initially occupied 6 patches and the two others occupied 5 patches in each landscape. We then followed community assembly for two weeks before forcing extinctions of all individuals of all species in selected patches and following the recolonization of those patches for two more weeks. Along with the landscapes, we also kept 9 monocultures (3 replicates per species) in single patches to provide a training data set for automated species identification (Pennekamp, Griffiths, et al., 2017).

#### Extinction patterns

The extinction patterns (Fig. S1) were chosen to either maximize (clustered extinctions) or minimize (dispersed extinctions) the percentage of like adjacencies (PLADJ). The PLADJ is calculated as the proportion of connections in a landscape that link two patches of the same kind (i.e., perturbed with perturbed or unperturbed with unperturbed) and is a measure of the spatial autocorrelation of the extinctions (PLADJ is close to 1 when extinctions are clustered, and close to 0 when they are dispersed). Because the landscapes are relatively small, the connectivity (i.e., the number of connections) of a patch varies depending on their position in the landscape. In order to minimize potential edge effects, we chose to draw the perturbed patches only from the sets of patches with a mean connectivity of three, which is the mean connectivity of the landscape. This ensured that corners, edges and central patches were equally represented in clustered and dispersed treatments, making them similar in terms of position relative to the edge. The drawing of extinction patterns was done by *i)* calculating the mean connectivity of all sets of 4 or 8 patches and keeping only those of connectivity 3, *ii)* calculating the PLADJ of the remaining sets and keeping only those with the highest PLADJ (for clustered extinctions) or lowest PLADJ (for dispersed extinctions) and *iii)* drawing an extinction pattern for each landscape among the remaining sets. We performed local patch extinctions by transferring the content of unperturbed patches to an identical new landscape in which perturbed patches were not transferred and replaced by fresh bacterized medium instead.

#### Data acquisition

The 2 mL of medium taken out of the patches and monocultures during medium replacement were used as samples to estimate the density of each species in each patch. For each patch and monoculture, 250 μL were put between two microscope slides (height: 500 μm) and filmed using an optical stereo-microscope (Perfex Pro 10) coupled with a camera (Perfex SC38800) for 10 seconds (150 frames).

#### Species identification

The three species differ in size, shape and behavior which allows for automated species identification (Pennekamp, Griffiths, et al., 2017). The videos were analyzed with the Bemovi R-package (version 1.0) (Pennekamp, Schtickzelle, et al., 2015) to track individuals and characterize their shape and trajectories (speed, size). The individuals were then identified from their characteristics (entire output of bemovi analysis) using a random forest algorithm (R-package randomForest version 4.6-14) trained on videos of the monocultures filmed on the same day (Pennekamp, Griffiths, et al., 2017). We rejected all the individuals with an identification confidence (proportion of trees leading to that identification) lower than 0.8 as a good compromise between the number of observations discarded and the confidence of identification (Fig. S4).

#### Diversity measures

*α*-diversity was measured as the inverse of Simpson’s index, which represents an effective number of species (Jost, 2006), and takes the relative abundance of species into account. We used the function beta.div.comp (R-package adespatial version 0.3-8, Ruzicka-based index) to compute the total *β*-diversity among the patches of a landscape (Legendre and De Cáceres, 2013).

#### Statistical analyses

All statistical analyses were conducted in R (version 4.0.2). To test the relative effects of spatial autocorrelation and amount of local extinctions on metacommunitiy properties, we studied 4 metrics (biomass, *α*-diversity, *β*-diversity and biomass recovery time) using mixed-effects models (R-package lme4 version 1.1-23) with measurement point and landscape ID (for patch level metrics) as random effects to account for non-independence of measures taken the same day and measures taken within one landscape. Fixed effects were the autocorrelation of extinctions, the amount of extinctions, as well as their interaction. Response variables were normalized using the R-package bestNormalize (version 1.6.1). The biomass in each patch was estimated using the bioarea per volume, a measure of the total surface of organisms visible in a video divided by the volume of medium in the camera field. The biomass recovery from extinction was estimated as the time needed to reach a bioarea per volume higher that the 2.5% quantile of pre-extinction bioarea in a given patch. This time span is hereafter referred to as recovery time.

For each statistical model, we performed AICc-based model selection on all models from the intercept to the full model. We used the weighted average of the model selection for predictions.

The direct effects of extinctions (*i.e.*, the variations of biomass and *α*-diversity in perturbed patches as well as the variations of *β*-diversity; Fig. 1) were estimated using all the measurements obtained in perturbed patches in the two weeks following the extinctions. We expected the indirect effects of extinctions (*i.e.*, the variations of biomass and *α*-diversity in unperturbed patches; Fig. 3) to be much more elusive, so we used only the data from unperturbed patches directly adjacent to perturbed patches. We expected indirect effects on biomass (*i.e.*, a reduction of the biomass of unperturbed patches due to reduced fluxes from perturbed patches) to happen early in the recolonization process, so we estimated them using only the data obtained just after the perturbations (from the two measurements following the extinctions, Fig. 3b). On the contrary, we expected indirect effects on *α*-diversity to happen late in the recolonization process (once the biomass in perturbed patches was high enough to have an effect on the composition of unperturbed patches) so we estimated them using data obtained near the end of the experiments (from the last two measurements made, Fig. 3a).

**Figure 1.**
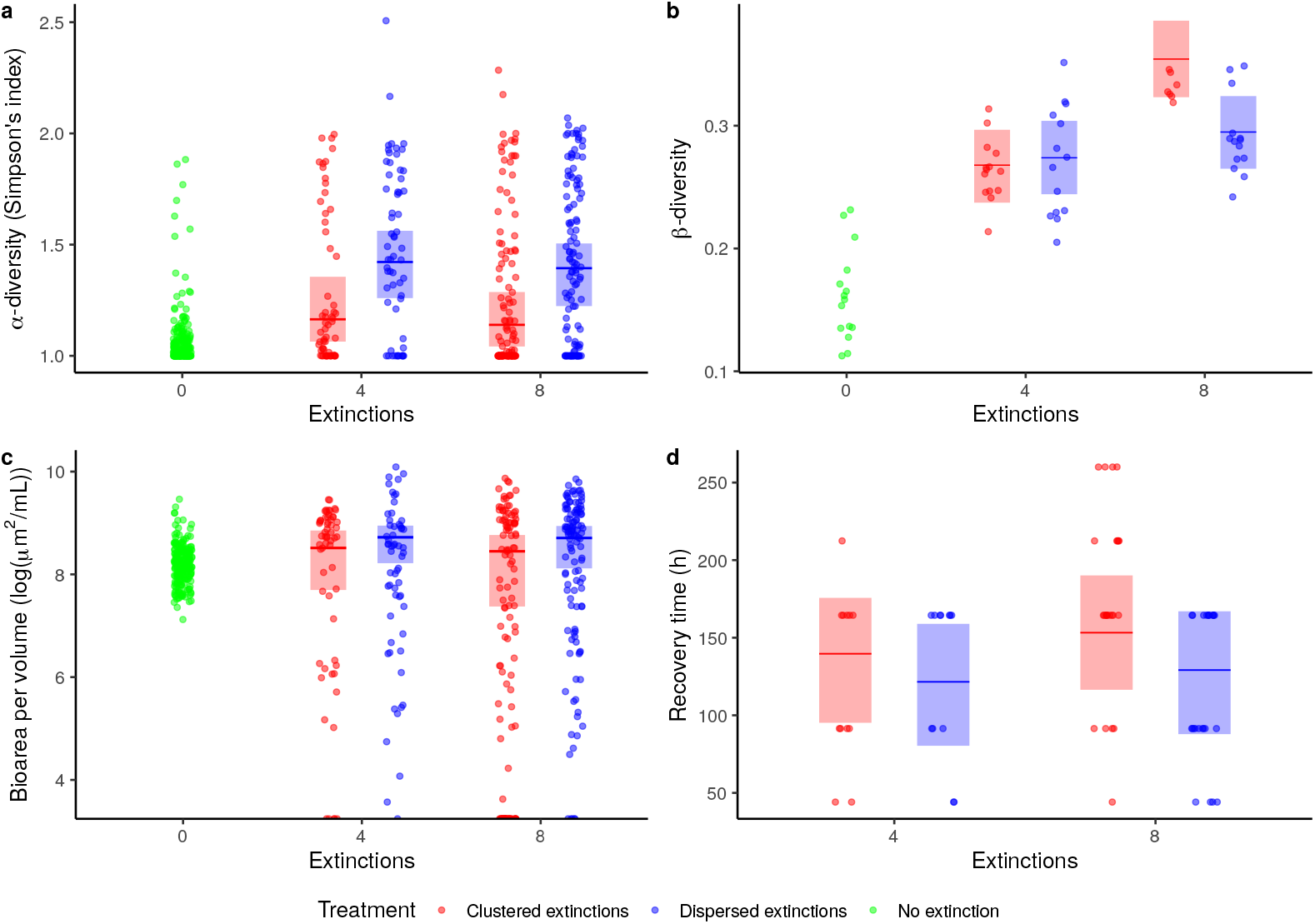
Observed response variables in the experiments (dots) and averaged mixed model predictions (medians and 95% confidence intervals; Tab. S3) from the extinction events to the end of the experiments. (a) *α*-diversity (measured as Simpson’s index) in perturbed patches (blue, red) and patches from landscapes with no extinctions (green), (b) *β*-diversity in all landscapes, (c) Bioarea in perturbed patches and patches from landscapes with no extinctions and (d) biomass recovery time in perturbed patches.

### Metacommunity model

We developed a mathematical model describing the dynamics of a competitive metacommunity of *n* species characterized by demographic and interaction parameters in landscapes similar to those used experimentally (i.e., a square lattice of 4 by 4 patches). We used Bayesian inference of demographic parameters on times series from the experimental single-patch cultures to parameterize the model (see below for details). We simulated dynamics using the same extinction plans as in the microcosm experiments with 100 replicates for each treatment.

#### Metacommunity dynamics

We used a set of ordinary differential equations to describe the dynamics of metacommunities (Eq. 1), where the terms describe the local dynamics (*f*), the emigration (*g*) and the immigration (*h*) of species *i* in patch *k*, with *N_i,k_* as the density of species *i* in patch *k*.

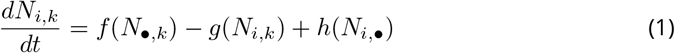

The local dynamics are described by a competitive Lotka-Volterra equation (Eq. 2) were *N_i,k_* grows logistically (*r_i_*: growth rate, *α_i,i_*: intraspecific competition) and is down-regulated by inter-specific competition (*α_i,j_*).

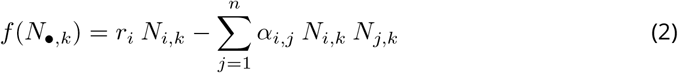

The number of individuals emigrating from a patch *k* is defined by a constant dispersal rate *m_i_* (Eq. 3).

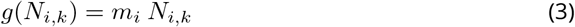

In analogy, we obtain the number of individuals immigrating into patch *k* as follows (Eq. 4):

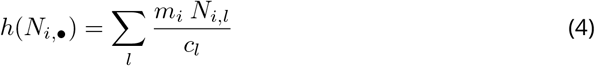

where *l* are the patches adjacent to *k* and *c_l_* is the number of connections leaving the patch *l*.

#### Parameterization of the model

We used four different sets of parameters (hereafter referred to as “scenarios of species interactions”) to investigate which processes may be responsible for the patterns observed experimentally. Two scenarios of species interactions (“empirical interactions” and “competition-colonization trade-off”) used demographic parameters (population growth rates *r_i_* and competition coefficients *α_i,j_*) fitted from empirical time series and were expected to most closely reproduce the experimental data. One scenario (“randomized interactions”) used the same competition coefficients but randomly shuffled between species in order to investigate whether the results were specific to our experimental community or if they could arise in other competitive communities with a different structure but similar overall interactions strength. The last scenario (“no interspecific interactions”) ignored interspecific interactions altogether and was thought of as a control scenario.

##### Empirical interactions

We parameterized the model using single-patch time series of mono-, bi- (cultures of *Blepharisma* sp. with *T. thermophila* and of *Blepharisma* sp. with *Colpidium* sp.) and tri-specific cultures from the experiments (three replicates of each culture). We fitted competitive Lotka-Volterra equations to the data using Bayesian inference (R-package Rstan version 2.19.3) (Feng et al., 2020; Rosenbaum et al., 2019). We fitted a single set of parameters (three *r_i_* and a 3 by 3 matrix of *α_i,j_*) over all replicates of all single-patch cultures (one curve per culture, with different initial conditions *N*_0_ for each culture), using lowly informative priors (Tab. S1) and assuming a negative binomial distribution of the residuals. We fit the model using the No U-Turn Sampler (NUTS) with three chains each of total length 10 000 (of which 2 000 steps were discarded as warm-up). We used default parameters for the sampler, except for the control parameters “adapt_delta” (set at 0.9) and “max_treedepth” (set at 12). The average fit can be found for visual inspection in Fig. S5.

This allowed us to infer values of population growth rates (*r_i_*) and competition coefficients (*α_i,j_*) for which the model yields dynamics that are quantitatively similar to the dynamics of the experimental community. We used the same dispersal rates for all three species (*m_i_* = 1/100).

##### Competition-colonization trade-off

We used the fitted values from the experimental results for the Lotka-Volterra parameters (*r_i_*, *α_i,j_*) and used different dispersal rates for each species (*m_i_* = {1/50, 1/100, 1/500}) with the most (resp. least) competitive species having the lowest (resp. highest) dispersal rate, resulting in a trade-off between competition and colonization.

##### Randomized interactions

We used the same parameters as in the “empirical interactions” scenario but we randomized interspecific interactions (i.e., the off-diagonal terms of the competition matrix: *α_i,j_, i* ≠ *j*). We randomly changed the position of the interaction terms while keeping each *α_i,j_* associated to the same *α_j,i_*.

##### No interspecific interactions

We used the same parameters as in the “empirical interactions” scenario but we set the interspecific interaction terms (*α_i,j_, i* ≠ *j*) to be zero. This results in a community where species do not experience interspecific competition. This scenario can be seen as a null model to investigate whether experimental results depended on interspecific interactions (in which case they should not be reproduced by this scenario) or whether they resulted from the neutral diffusion of species on a lattice (in which case they should be reproduced by this scenario).

#### Sensitivity analysis

We ran additional simulations to explore if our findings were robust to variations in landscape size and dispersal rates.

##### Landscape size

We ran the simulations on larger landscapes (a square lattice of 16 by 16 patches) with the same proportion of extinctions (either no extinctions, extinctions in a quarter of the patches or extinctions in half of the patches) (Fig. S7 and S8).

##### Dispersal rate

Finally, we ran simulations for larger (times 2 and times 5) and smaller (divided by 2 or 5) dispersal rates (Fig. S9 to S16).

## Results

### The role of the spatial distribution of extinctions

In the experiments, both local and regional effects of local patch extinctions were mainly determined by the spatial autocorrelation of extinctions. Except for *β*-diversity, the amount of extinctions alone only had a marginal effect on the outcome of the experiment as indicated by model selection (Fig. 1; Tab. S3). For the local variables studied (*α*-diversity, bioarea and bioarea recovery time), the autocorrelation of extinctions was found to be more important than the amount of extinctions (Tab. S3). Both *α*-diversity in unperturbed patches (Tab. S4b) and *β*-diversity (Tab. S3b) were mostly explained by the interaction between autocorrelation and amount of extinctions (statistical models without the interactions had either a null (for *β*-diversity) or low (for *α*-diversity) weight).

Numerical simulations of our metacommunity model with the same spatial configuration and extinctions patterns reproduced these results (a weak effect of the amount of extinctions compared to that of their spatial arrangement) for all competition scenarios (Fig. 2 and 4).

**Figure 2.**
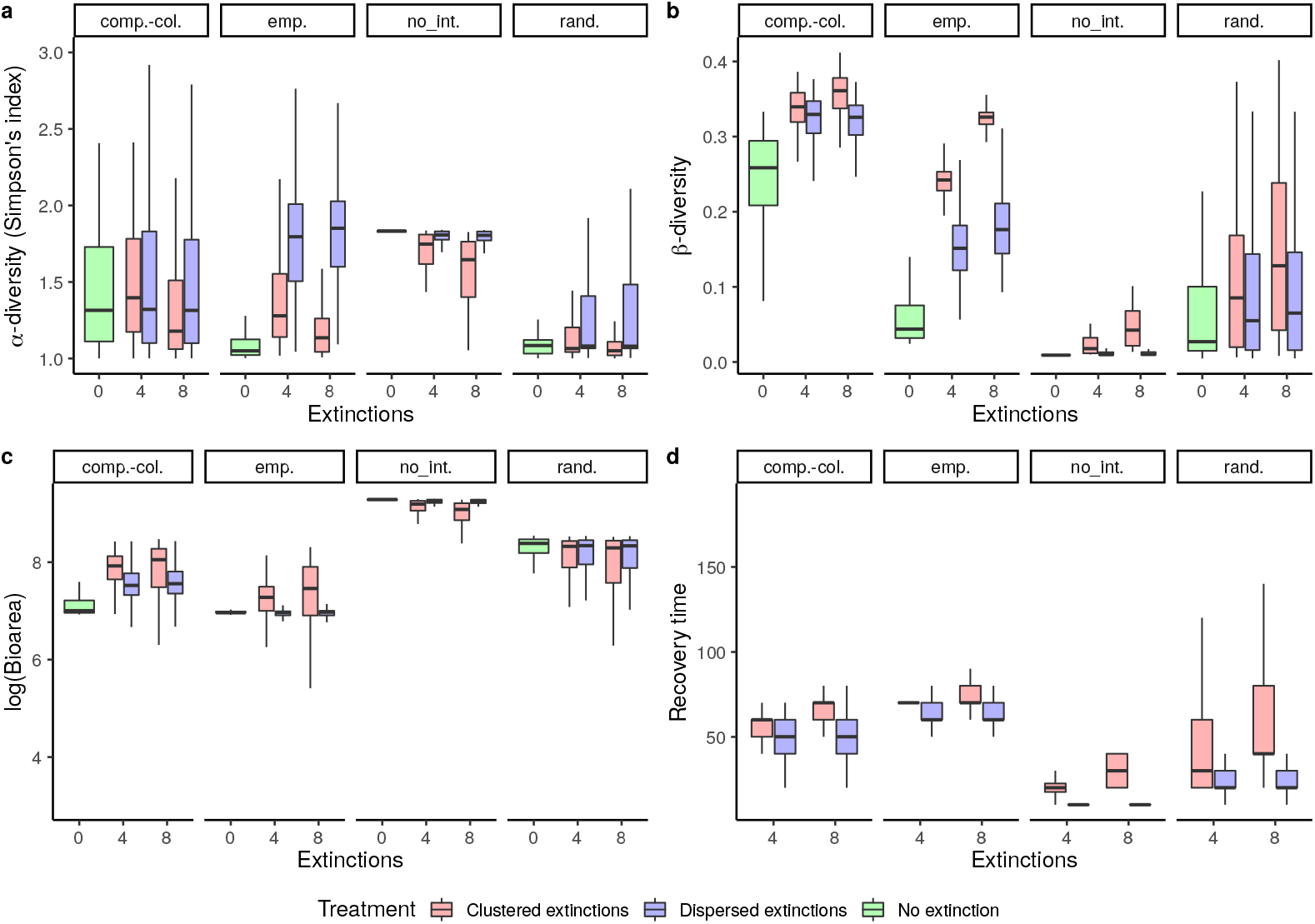
Observed response variables in numerical simulations of the metacommunity model displaying different metrics after the extinction events. (a) *α*-diversity (measured as Simpson’s index) in perturbed patches (blue, red) and patches from landscapes with no extinctions (green), (b) *β*-diversity in all landscapes, (c) Bioarea in perturbed patches and patches from landscapes with no extinctions and (d) biomass recovery time in perturbed patches. The top labels denote the scenarios of species interactions: “emp.” for “empirical interactions”, “comp.-col.” for “competition-colonization trade-off”, “rand.” for “randomized interactions” and “no int.” for “no interspecific interactions”.

### Direct effects — recolonization dynamics in perturbed patches

We first consider the recolonization dynamics of biomass and *α*-diversity in perturbed patches.

#### Biomass

The bioarea per volume, as proxi for biomass in a given patch, after local patch extinctions was slightly higher in perturbed patches from landscapes with dispersed extinctions than in landscapes with clustered extinctions (Fig. 1c, median predictions : ~ 6000 μm^2^ mL^−1^ vs. ~ 5000 μm^2^ mL^−1^). Note that this effect is weak as indicated by model selection which ranks the intercept model second with an AICc weight of 0.27 (Tab. S3). The recovery time needed to reach a bioarea higher than the 2.5% quantile of the pre-extinction bioarea was shorter in case of dispersed extinctions compared to clustered extinctions, and it slightly increased with the amount of extinctions (Tab. S3 and Fig. 1d and S18; median mixed model predictions: 4 dispersed: 122 h, 8 dispersed: 130 h, 4 clustered: 139 h, 8 clustered: 134 h).

In simulations of the metacommunity model, recovery times (Fig. 2d) depended greatly on the scenario of species interactions: it was shorter in the absence of interspecific interactions (scenario: “no interspecific interactions”) and with randomized interactions (“randomized interactions”), and longer for fitted interaction terms (“empirical interactions” and “competition-colonization trade-off”). However, the differences between treatments were qualitatively similar between all interaction scenarios: the recovery times were shorter for dispersed extinctions than for clustered extinctions. In landscapes with dispersed extinctions, the recovery times were not affected by the amount of extinctions. By contrast, in landscapes with clustered extinctions, the recovery times increased with the amount of extinctions. It is noteworthy that, in general, the recovery times were much shorter (less than 100 time units) than what we found experimentally, probably because dispersal in the experiments happened over discrete time interval (4 h periods, three times per week) resulting in a lag in recolonization dynamics.

#### *α*-diversity

In patches from control landscapes (i.e., landscapes without any patch extinctions), *α*-diversity increased at first as species dispersed between patches but quickly fell to 1 (the minimal value) as *Blepharisma* sp. finally excluded the two other species and dominated the community (Fig. S6). In perturbed patches of the landscapes with extinction treatments, *α*-diversity was higher during the recolonization process in comparison to the control landscapes since the species were present in more even densities (Fig. 1a and S6). This effect was stronger for dispersed extinctions than for clustered extinctions (Fig. 1a).

In simulations from the metacommunity model, *α*-diversity patterns depended on the scenario of species interactions (Fig. 2a). In the absence of interspecific interactions (“no inter-specific interactions”), the three species could coexist locally and the *α*-diversity stayed high in patches from control landscapes. In perturbed patches, the *α*-diversity was 1 right after extinction but quickly came back to pre-extinction levels as all species recolonized (Fig. S17). This recovery was faster for dispersed than for clustered extinctions and in landscapes with 4 rather than 8 extinctions. In all three other scenarios (“empirical interactions”, “randomized interactions” and “competition-colonization trade-off”), interspecific interactions resulted in competitive exclusion. As a consequence, *α*-diversity was fairly low in control landscapes (Fig. 2a). In the perturbed patches of the landscapes with extinction treatments, *α*-diversity during the recolonisation process was higher (for all treatments) than in the patches from control landscapes. *α*-diversity was highly variable in time during the recolonization process (Fig. S17). In all scenarios, *α*-diversity in patches from dispersed extinction treatments was higher early in the recolonization process but then decreased quickly. Later in the recolonisation process, *α*-diversity was higher in patches from clustered extinction treatments than in patches from dispersed extinction treatments.

### Indirect effects — spread of extinctions effects to unperturbed patches and at the regional scale

As local events can spread in space and have regional consequences, we now focus on the indirect effects of local patch extinctions on undisturbed patches (biomass and *α*-diversity) and on regional effects (*β*-diversity).

#### Biomass

We observed no strong difference in bioarea per volume between treatments (Fig. 3b and 4b). Although the bioarea predictions from the mixed model are slightly higher in unperturbed patches than in patches from control landscapes, both empirical data and the statistical models predictions are largely overlapping between treatments.

**Figure 3.**
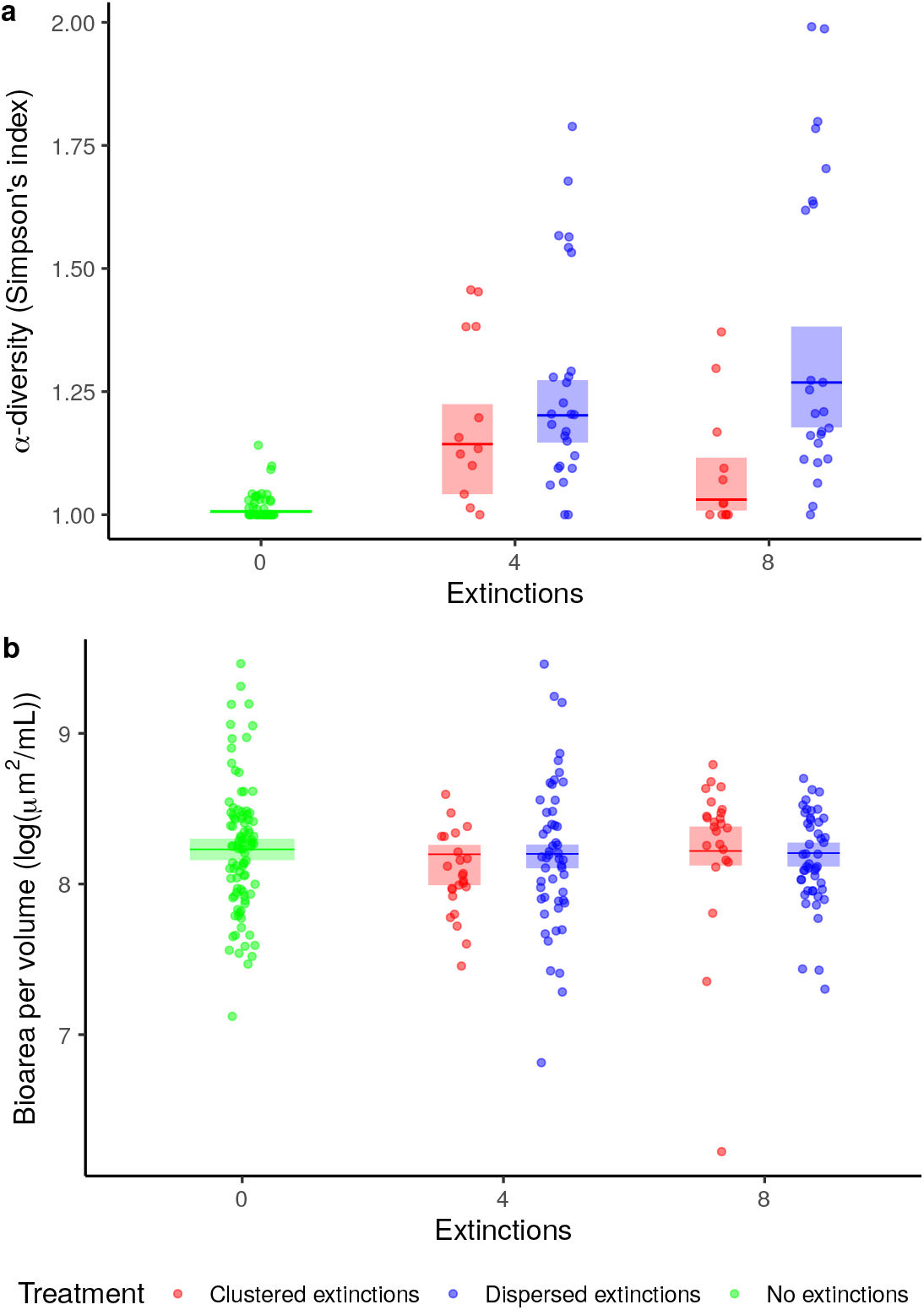
Observed response variables in the experiments (dots) and averaged mixed model predictions (medians and 95% confidence intervals; Tab. S4) in unperturbed patches adjacent to at least one perturbed patch (blue, red) and in control landscapes (green). (a) *α*-diversity (measured as Simpson’s index) in unperturbed patches at the last two measurements, (b) bioarea in unperturbed patches (for the two measurements following the extinctions).

**Figure 4.**
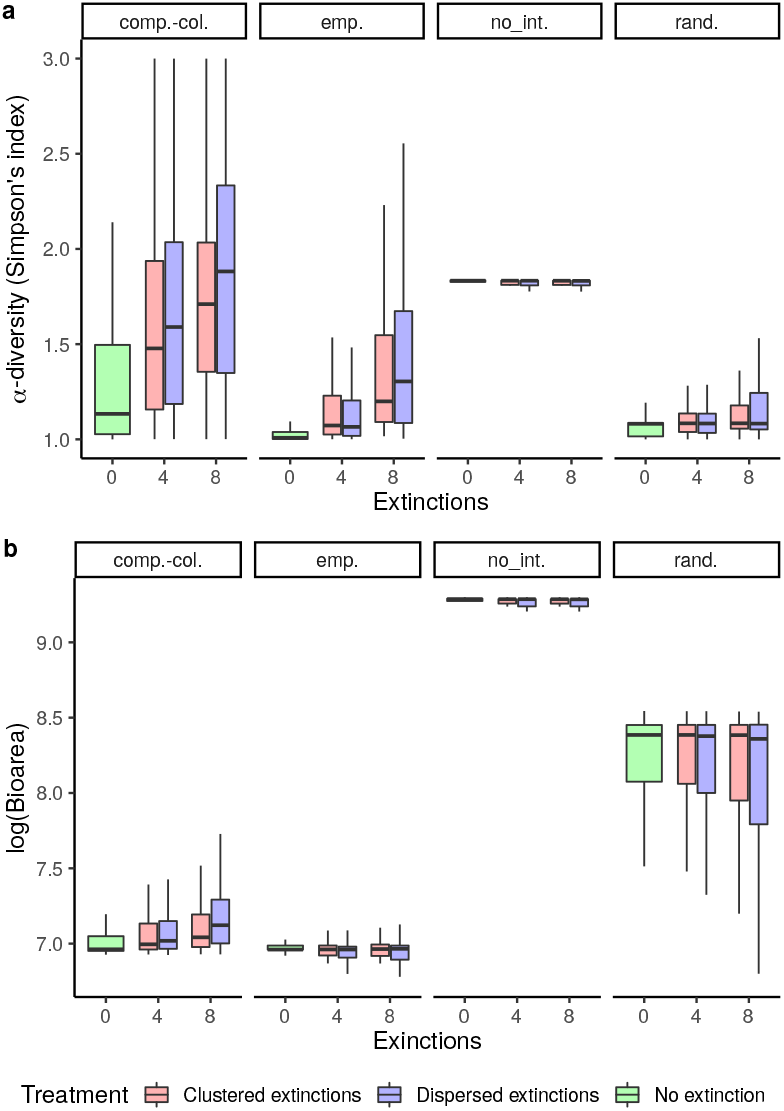
Observed response variables in numerical simulations of the metacommunity model showing *α*-diversity (measured as Simpson’s index) (a) and biomass (b) in unperturbed patches adjacent to at least one perturbed patches (blue, red) and in control landscapes (green) after extinction events. The top labels denote the scenarios of species interactions: “emp.” for “empirical interactions”, “comp.-col.” for “competition-colonization trade-off”, “rand.” for “randomized interactions” and “no int.” for “no interspecific interactions”.

#### *α*-diversity

Experimentally, *α*-diversity was higher in unperturbed patches than in patches from control landscapes, particularly for dispersed extinctions (Fig. 3a). Most of the variation between treatments was explained by the spatial autocorrelation of extinctions rather than the amount of extinctions (Tab. S4b). Interestingly, the effect of the amount of extinctions depended on their spatial organization: under clustered extinctions, the *α*-diversity in unperturbed patches decreased with the amount of extinctions but it increased under dispersed extinctions (Fig. 3a).

The results from the simulations of the metacommunity model depended on the scenarios of species interactions (Fig. 4a): in the absence of interspecific competition (“no interspecific interactions”), *α*-diversity levels were similar in unperturbed patches (across all treatments) and patches from control landscapes. In every other scenario (“empirical interactions”, “randomized interactions” and “competition-colonization trade-off”), *α*-diversity was higher in unperturbed patches than in patches from control landscapes. In line with experimental results, *α*-diversity was higher for treatments with dispersed extinctions. *α*-diversity also increased with the amount of extinctions. Although these results were qualitatively similar across the scenarios that included interspecific competition (“randomized interactions”, “empirical interactions” and “competition-colonization trade-off”), the effect sizes were highly variable: empirical interactions yielded effect sizes consistent with the experimental results (according to qualitative visual inspection), while randomized interactions yielded smaller effects and the “competition-colonization trade-off” scenario yielded stronger effects.

#### *β*-diversity

In control landscapes, *β*-diversity was fairly low because the patches ended up being homogeneous and dominated by *Blepharisma* sp. (Fig. S6). *β*-diversity was higher in landscapes with extinctions than in control landscapes because of differences in species composition and density between perturbed and unperturbed patches (Fig. S6). This effect was stronger for 8 extinctions than for 4 extinctions, particularly for clustered extinctions (Fig. 1b).

In simulations of the metacommunity model, these results held qualitatively for all competition scenarios (Fig. 2b): *β*-diversity was higher in landscapes with extinctions than in control landscapes. Among landscapes with extinctions, *β*-diversity generally increased with spatial autocorrelation and amount of extinctions. These effects were strong and on par with experimental effect sizes for realistic interaction matrices (scenarios “empirical interactions” and “competition-colonization trade-off”). They were weaker for randomized interaction matrices (“randomized interactions” scenario) and negligible in the absence of interspecific interactions (“no interspecific interactions” scenario).

### Sensitivity to landscape size and dispersal rates

The simulations on larger landscapes (16*16 patches) yielded results (Fig. S7 and S8) remarkably consistent with those discussed above (landscapes of 4*4 patches, Fig. 2 and 4). Our results were more sensitive to dispersal rates, but most qualitative patterns described for the “empirical interactions” and “competition-colonization trade-off” scenarios (e.g., stronger influence of the spatial autocorrelation than the amount of extinctions, higher *β*-diversity for clustered extinctions, higher *α*-diversity spillover and faster biomass recovery for dispersed extinctions) were coherent for dispersal rates up to 2 times stronger/weaker than our standard simulations (Fig. S9 to S16).

## Discussion

### The role of the spatial distribution of the extinctions

Our work clearly shows that recovery from extinctions depends more on the spatial features of local patch extinctions (such as the connectivity between perturbed and unperturbed patches) than on interspecific interactions or on the amount of patches affected. More specifically, our experiments clearly showed that the spatial autocorrelation of extinctions had stronger effects than the amount of extinctions per se on all metacommunity metrics measured, including biomass, recovery time, *α*- and *β*-diversity (Tab. S3). These empirical findings were confirmed by our theoretical model, regardless of the specific scenario. The main factor driving these results can be linked to the connectivity and distance between perturbed and unperturbed patches: in the dispersed extinction treatments, perturbed patches were closer and better connected to unperturbed patches than in the clustered extinction treatments (Tab. S2; Fig. S18 and S19), which modulated the recovery dynamics. These results can be interpreted as differences in recovery regimes across spatial treatments: clustered extinctions, characterized by a weak connectivity between perturbed and unperturbed patches, result in what Zelnik et al. (2019) described as a “rescue recovery regime”, while dispersed extinctions, characterized by a strong connectivity between perturbed and unperturbed patches, result in a “mixing recovery regime”. Under the “rescue” regime, dispersal between perturbed and unperturbed patches is marginal compared to local dynamics. Perturbed and unperturbed patches are strongly differentiated, and the recovery dynamics mainly rely on local growth. Because of this strong differentiation, *β*-diversity was higher than in the “clustered extinctions” treatment, but the high *α*-diversity of perturbed patches did not spill over much to unperturbed patches. Under the “mixing” regime, dispersal between perturbed and unperturbed patches is on par with local dynamics. Perturbed and unperturbed patches are well mixed, and both local growth and dispersal from perturbed patches participate substantially to the recovery. Because of the mixing between perturbed and unperturbed patches, *α*-diversity in the “dispersed extinctions” treatment in unperturbed patches increased greatly (due to dispersal from perturbed patches), but *β*-diversity was lower than in the “clustered extinctions” treatment.

#### Influence of distance and connectivity to unperturbed patches

Statistically determining how the local properties of a given perturbed patch (namely the distance to the closest unperturbed patch and the number of adjacent unperturbed patches) affect its recovery is difficult from our experimental data because of the low variability and redundancy of these indicators (Fig. S18 and S19). However, the analysis of the simulations of large landscapes (Fig. S20, S21 and S22) gives us a hint about the underlying mechanisms. Overall, the recovery dynamics in a given patch seemed to be mainly determined by the distance to the closest unperturbed patches (which is directly related to the size of the extinction cluster).

In terms of biomass, the recovery time of perturbed patches increased linearly with the distance to the closest unperturbed patch (Fig. S20a) but was mainly unaffected by the connectivity to unperturbed patches (number of adjacent unperturbed patches, Fig. S20b). This was in accordance with the experimental recovery dynamics (Fig. S18) where the patches further away from unperturbed patches recovered more slowly.

The local diversity of perturbed patches was also mainly related to the distance to the closest unperturbed patches (Fig. S21): *α*-diversity was the highest in perturbed patches directly adjacent to unperturbed patches (distance = 1) and decreased with the distance to unperturbed patches (Fig. S21a), because patches far away from unperturbed patches were either not recolonized or recolonized only by the better disperser. The connectivity to unperturbed patches hardly affected *α*-diversity, as all patches adjacent to at least one unperturbed patch (connectivity greater or equal to 1, Fig. S21b) had similar *α*-diversity. This is once again coherent with experimental results: perturbed patches from the dispersed treatments were all adjacent to unperturbed patches and had a high *α*-diversity, while perturbed patches from the clustered treatments were further away from unperturbed patches and had a lower *α*-diversity (Fig. S19).

Lastly, *β*-diversity was also determined by connectivity and distance between perturbed and unperturbed patches: the *β*-diversity of a landscape increased with the average distance between perturbed and unperturbed patches and decreased with the average connectivity between perturbed and unperturbed patches (Fig. S22).

These quantitative effects of the distance between perturbed and unperturbed patches explain the differences between dispersed extinctions (that result in perturbed patches being mainly adjacent to unperturbed patches) and clustered extinctions (that result in a greater distance between perturbed and unperturbed patches). It also explains why the amount of extinctions usually had a marginal effect in dispersed treatments compared to clustered treatments (Fig. 1 and 2): increasing the amount of extinctions did not increase the distance from perturbed to unperturbed patches for dispersed extinctions (Tab. S2). On the contrary, more clustered extinctions resulted in larger clusters and thus in a greater distance from perturbed to unperturbed patches (Tab. S2).

### Direct effects of extinctions

#### Biomass recovery

Experimental data and simulations support the conclusion that simultaneously increasing the rate and autocorrelation of extinctions increases the time needed for a metacommunity to recover its pre-extinction biomass (Fig. 1d and 2d). These results were surprisingly consistent between the experiments and the various simulations scenarios, highlighting that this pattern does not depend on species interactions but rather on the geometry of the patches to be recolonized. A high amount of spatially clustered extinctions increases the recovery time by creating large areas of perturbed patches, thus increasing the average distance and reducing the average connectivity between perturbed and unperturbed patches (Tab. S2). As above, this can be discussed from a recovery regime perspective (Zelnik et al., 2019): dispersed extinctions result in a “mixing recovery regime” where perturbed and unperturbed patches are well mixed and dispersal, in combination with local population growth, qualitatively participates to biomass recovery. Clustered extinctions result in a “rescue recovery regime” where biomass recovery relies mainly on local population growth and is thus slower.

Additionally, both experimentally and in model simulations, perturbed patches had a slightly higher biomass after recovery than patches from unperturbed landscapes (Fig. 1c and 2c). This is because unperturbed patches mainly had the better competitor left (*Blepharisma* sp., Fig. S6), while all three species persisted in perturbed patches. Since poorly competitive species (especially *Colpidium* sp.) reached a higher biomass than *Blepharisma* sp., perturbed patches had a higher biomass. This result should hold for communities dominated by highly competitive but slowly reproducing species that do not reach high densities (e.g., if there is a trade-off between population growth rate and competitive ability rather than the often assumed trade-off between population growth rate and carrying capacity; for a discussion, see Mallet 2012) or when populations are able to overshoot their equilibrium density. This should however not be the case for communities where the dominant species happens to reach higher equilibrium densities, as it is the case in forests, for instance, where transiently recolonising species (e.g., grasses or shrubs) do not accumulate biomass and are slowly replaced by dominant species that do (trees).

#### *α*-diversity

Local patch extinctions generally increased *α*-diversity: experimentally, unperturbed patches reached a state where *Blepharisma* sp. was largely dominant, sometimes to the point where *T. thermophila* and *Colpidium* sp. were locally excluded. In control landscapes, this resulted in the extinction of *T. thermophila* at the landscape scale. As a result, *α*-diversity was low in control landscapes and in unperturbed patches (Fig. 3a). In perturbed patches, all three species persisted during the recolonization process, resulting in higher *α*-diversity (Fig. 1a) compared to unperturbed patches from the same landscapes or from control landscapes (Fig. 3a). This result was also observed in all simulations of the metacommunity model, except in the absence of interspecific competition (“no interspecific interactions” scenario) since no competitive exclusion occurs in that case (Fig. 2a). The persistence of less competitive species in perturbed patches during the recolonisation process can be explained both by the decrease in population density and by a competition-colonization trade-off across the three species: the low population density after extinction events decreases the intensity of competition, while the competition-colonization trade-off delays the recolonization by *Blepharisma* sp., both processes resulting in the delay of competitive exclusion. Since the increased *α*-diversity was observed in simulations without a competition-colonization trade-off (i.e., scenarios “randomized interactions” and “empirical interactions”; Fig. 2a), such a trade-off is not necessary for local extinctions to increase *α*-diversity, even though the trade-off increased *α*-diversity even more. These results are similar to the effect described in the intermediate disturbance hypothesis which predicts that some degree of perturbation should result in a higher local and regional biodiversity by reducing the abundance of competitively dominant species and allowing the persistence of early succesional species (Shea et al., 2004; Wilkinson, 1999). However, previous experiments on similar systems found that local patch extinctions decreased local diversity (Cadotte, 2007). This can be explained by differences in metacommunity composition: metacommunities skewed towards early-succesional species should exhibit the *α*-diversity increase observed here, while metacommunities skewed towards late-succesional species (as in Cadotte 2007) should see *α*-diversity decrease with local patch extinctions.

Clearly, these effects may be relevant in the context of ecosystem management: while local perturbations (here in their most extreme form, the extinction of all species) decrease biomass, they can also allow the persistence of species that would otherwise be excluded and lead to an increased local diversity.

### Indirect effects

Besides the direct effects discussed above, local patch extinctions may also have indirect effects at the regional scale by altering species densities and composition in unperturbed patches (Gilarranz et al., 2017; Zelnik et al., 2019).

#### Biomass

Biomass in unperturbed patches was mainly unaffected by local patch extinctions: biomass distributions largely overlapped between unperturbed patches and patches from control landscapes (experimentally: Fig. 3b; in simualtions: Fig. 4b). Despite reduced fluxes from perturbed patches, the density in unperturbed patches did not decrease. This can be explained by local dynamics (population growth) being faster than spatial dynamics (dispersal). In this case, the adverse effect of local extinctions (decreased biomass) does not spread to unperturbed patches. However, in metacommunities with strong dispersal, unperturbed patches should also experience reduced biomass. While we did not observe a decrease of biomass in unperturbed patches, probably because local dynamics were too fast for spatial dynamics to have an effect on these patches, previous theoretical work predicts that a local biomass reduction could spread in space if dispersal rates were high enough (Zelnik et al., 2019).

#### *α*-diversity

Experimentally, unperturbed patches in landscapes with extinctions were not dominated by *Blepharisma* sp. This is because dispersal of *T. thermophila* and *Colpidium* sp. from perturbed patches, where they were present in high density during the recolonization process, allowed these species to persist in unperturbed patches (Fig. S6). Their persistence increased *α*-diversity in unperturbed patches compared to patches from control landscapes that were mainly monospecific (Fig. 3a and S6). The increase of *α*-diversity was stronger in unperturbed patches from dispersed extinction treatments, as these patches were connected to more perturbed patches and thus received an increased amount of less competitive dispersers than unperturbed patches from clustered extinction treatments.

The increase of *α*-diversity following extinctions did not occur in the metacommunity model in the absence of interspecific competition (Fig. 4a; scenario “no interspecific interactions”), because competitive exclusion did not occur and therefore all three species were present in all patches. However, the patterns observed experimentally were recovered in all simulations that incorporated interspecific competition (Fig. 4a; scenarios “randomized interactions”, “empirical interactions” and “competition-colonization trade-off”), showing that local diversity maintenance by local extinctions is not restricted to our particular experimental community but can occur as long a some species excludes others.

It is worth noting that the increase in *α*-diversity was only observed in patches adjacent to perturbed patches, which could be described as an edge effect (in the sense that indirect effects are only observed at the edge of perturbed patches). This means that isolated extinction events don’t have large scale effects in our setting, as perturbed patches only have an effect on their local neighbourhood. Indirect effects, however, can affect large proportions of the landscape if extinctions are numerous and spatially dispersed (*e.g.*, in the treatment with eight dispersed extinctions, all eight unperturbed patches were adjacent to perturbed patches vs. only four in the eight clustered extinctions treatment). Dispersed extinctions thus have both a stronger effect on unperturbed patches and affect a greater number of unperturbed patches.

#### *β*-diversity

*β*-diversity was higher in landscapes that experienced local patch extinctions in comparison to control landscapes, both in experiments and in simulations including interspecific competitition (Fig. 1b and 2b). In the simulations without interspecific competition (Fig. 2b; scenario “no interspecific interactions”), *β*-diversity increased only marginally because all three species quickly recolonized the patches in the same proportion as in unperturbed patches. The increase in *β*-diversity following local patch extinctions (in experiments and in simulations with interspecific competition) can be explained by the fact that perturbed patches had a different species composition than unperturbed patches. In unperturbed patches communities were mainly composed of *Blepharisma* sp., while perturbed patches allowed for less competitive species to persist during the recolonization process. While we find a strictly increasing relationship between the amount of extinctions and *β*-diversity (Fig. 1b and 2b), Cadotte (2007) found a unimodal relationship between *β*-diversity and local patch extinction rates. While this seems contradictory, it is also possible that we did not cover enough amount of extinctions to uncover a unimodal relationship, as *β*-diversity could decrease when extinctions affect more patches.

By crossing the amount of extinctions and spatial autocorrelation treatments, we were able to show that the relationship between *β*-diversity and local patch extinctions is strongly dependant on the spatial distribution of extinctions: the increase in *β*-diversity was higher when extinctions were clustered than when they were dispersed in space. When extinctions were clustered, the connectivity between perturbed and unperturbed patches was fairly low, resulting in a strong differentiation between perturbed and unperturbed patches. When extinctions were dispersed, perturbed and unperturbed patches were well connected, resulting in a stronger mixing of communities between patches and a lower *β*-diversity.

### Perspectives

Clearly, we have used a number of simplifying assumptions in our metacommunity model as well as in the experimental work that could provide some interesting directions for future research.

Firstly, we consider only competitive interactions between species while natural communities consist of more diversified interactions, including predation, mutualism and parasitism, for example (Kéfi, Berlow, Wieters, Joppa, et al., 2015; Kéfi, Berlow, Wieters, Navarrete, et al., 2012). These interactions could complicate the response (Kéfi, Miele, et al., 2016) and affect the consequences of extinctions on ecological communities. Moreover, the sensitivity of species to local extinctions could depend on their trophic level, as demonstrated for habitat destruction (Liao et al., 2017; Ryser et al., 2019): top predators (or parasites) could be more vulnerable as they suffer both from the perturbation and from the reduction of their prey (or host) density. Specialized predators and parasites may also take longer to recolonize since they cannot return to perturbed patches while their prey (or host) is not present at a high enough density. Vice versa, other species could benefit from local extinctions through decreased predator or parasite pressures.

Secondly, the temporal scale of our study is very narrow as we consider a single event of synchronous extinctions. In nature, extinction events can potentially be asynchronous and recurring over time. Both the degree of synchrony and the frequency of extinction events could shape their consequences on metacommunity dynamics. A first intuitive approach to explore these directions would be to use a space-for-time substitution, and to consider the amount of extinctions (in space) as analogous to a frequency of extinctions (in time) and the spatial autocorrelation as analogous to the synchrony of extinctions. However, adding a temporal dimension could also lead to consequences unforseen in our mostly spatial setting, such as the synchrony/asynchrony of extinctions affecting metacommunity stability by affecting the synchrony/asynchrony of local community dynamics (Fox et al., 2017). Exploring these questions would thus require to go beyond a simple space-for-time substitution and to conduct new experiments on a larger temporal scale.

Thirdly, we ignore evolutionary processes although natural populations can readily adapt to environmental change. Increased amounts of local patch extinctions should select for higher dispersal rates (Bowler and Benton, 2005; Ronce, 2007), but increased spatial autocorrelation of extinctions could select for lower dispersal rates and longer dispersal distances (Fronhofer, Stelz, et al., 2014), which could result in opposite selective pressures if both increase at the same time. This could have implications for the dynamics of biodiversity because dispersal can mediate species coexistence (Hanski, 1983), diversity patterns (Laroche et al., 2016) and speciation (Pellissier, 2015). In particular, increased dispersal could synchronize metacommunities, making them more prone to global extinctions. Metacommunity synchrony could also be increased by the increasing spatial synchrony of climatic events (Di Cecco and Gouhier, 2018), as observed in the metapopulation of *Melitaea cinxia* (Kahilainen et al., 2018). On the other hand, evolutionary rescue could buffer the effects of disturbances, allowing metacommunities to persist in increasingly harsher environments (Bell and Gonzalez, 2011).

## Conclusion

Overall, our study shows that the effects of local patch extinctions in metacommunities strongly depend on the spatial distributions of extinctions. Local patch extinctions can increase both *α*-diversity and *β*-diversity by allowing weak competitors to persist in the metacommunity and by forcing a differentiation between perturbed and unperturbed patches.

Dispersal and connectivity between patches are central to recovery as they allow the recolonization of perturbed patches but also a mixing between perturbed and unperturbed patches, which can result in the spread of local extinction effects to unperturbed patches. In our setting, this spread was characterised by an increase in *α*-diversity in unperturbed patches through dispersal from species-rich, previously perturbed patches to species poor, unperturbed patches.

By determining the connectivity between perturbed and unperturbed patches, the spatial autocorrelation of extinctions modulates the dynamics after the extinction events: when extinctions are clustered, perturbed and unperturbed patches are weakly connected. This results in a slower biomass recovery, a weak spread of *α*-diversity and a high *β*-diversity as perturbed and unperturbed patches are differentiated. On the contrary, dispersed extinctions imply higher connectivity between perturbed and unperturbed patches which translates into a faster biomass recovery, a stronger spread of *α*-diversity and a lower *β*-diversity as perturbed and unperturbed patches are better mixed.

Our highly controlled experiment in combination with the theoretical model provide a proof-of-concept for the importance of taking into account the spatial distribution of disturbances in biodiversity research. Of course, applying our findings to specific, real-world ecosystems will require a combination of field data and system-specific models to better estimate the effects of local extinctions in more realistic settings. Nevertheless, our work highlights the importance of the spatial distribution of local extinctions when doing so.

## Author contributions

C.S., S.K. and E.A.F. conceived the study. C.S. and C.G.B. conducted the experiments. C.S. performed the statistical analyses. C.S., B.R. and E.A.F. performed the model fitting. C.S. analysed the mathematical model. C.S., S.K. and E.A.F. wrote the manuscript and all authors commented on the draft.

## Acknowledgements

The study was funded by a grant of the ENS to C.S. and CNRS funds to E.A.F. B.R. acknowledges the support of iDiv funded by the German Research Foundation (DFG–FZT 118, 202548816). Version 4 of this preprint has been peer-reviewed and recommended by Peer Community In Ecology (https://doi.org/10.24072/pci.ecology.100084). This is publication ISEM-2021-119 of the Institut des Sciences de l’Evolution – Montpellier.

## Data accessibility

Data and code are available on GitHub via Zenodo: https://doi.org/10.5281/zenodo.4660016

## Conflict of interest disclosure

The authors of this article declare that they have no financial conflict of interest with the content of this article. Emanuel A. Fronhofer is one of the PCI Ecology recommenders.

## Supplementary Material

### Supplementary Figures

**Figure S1.**
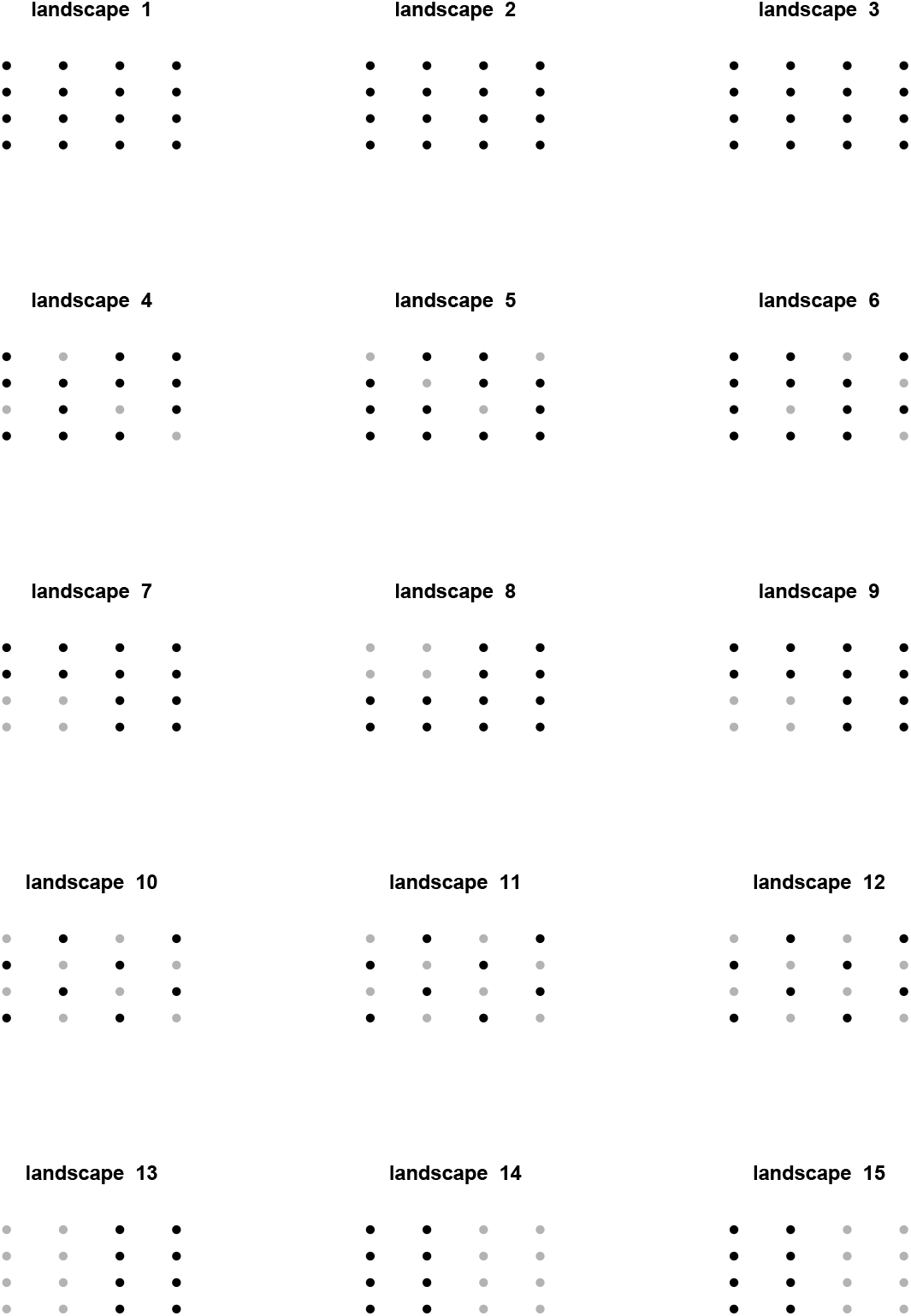
Positions of the extinctions (grey) in each landscape in the experimental setting. Landscapes 1-3: no extinction, landscapes 4-6: 4 dispersed extinctions, landscapes 7-8: 4 clustered extinctions, landscapes 10-12: 8 dispersed extinctions, landscapes 13-15: 8 clustered extinctions.

**Figure S2.**
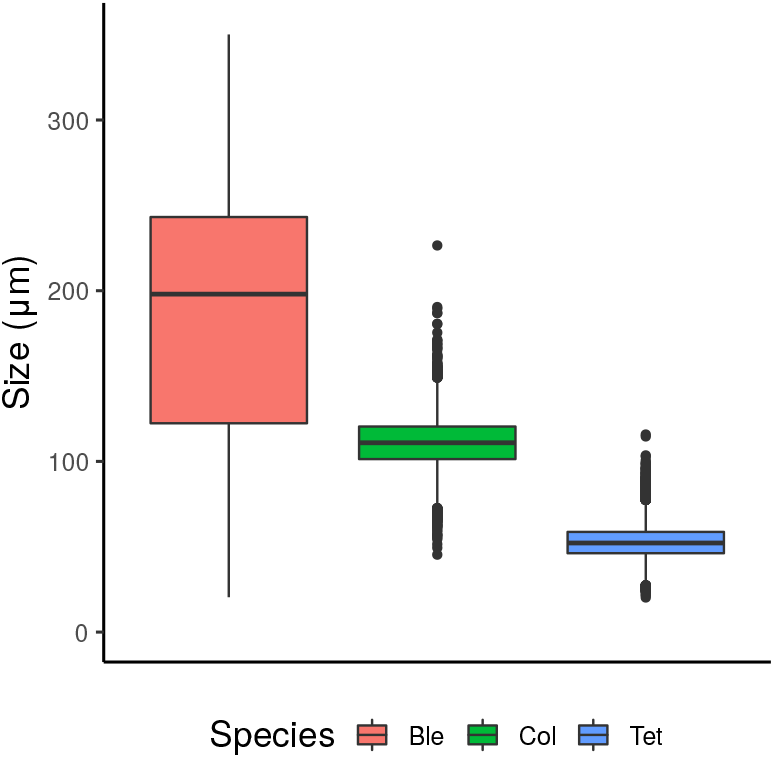
Size distributions of *T. thermophila* (Tet), *Colpidium* sp. (Col) and *Blepharisma* sp. (Ble) in monocultures.

**Figure S3.**
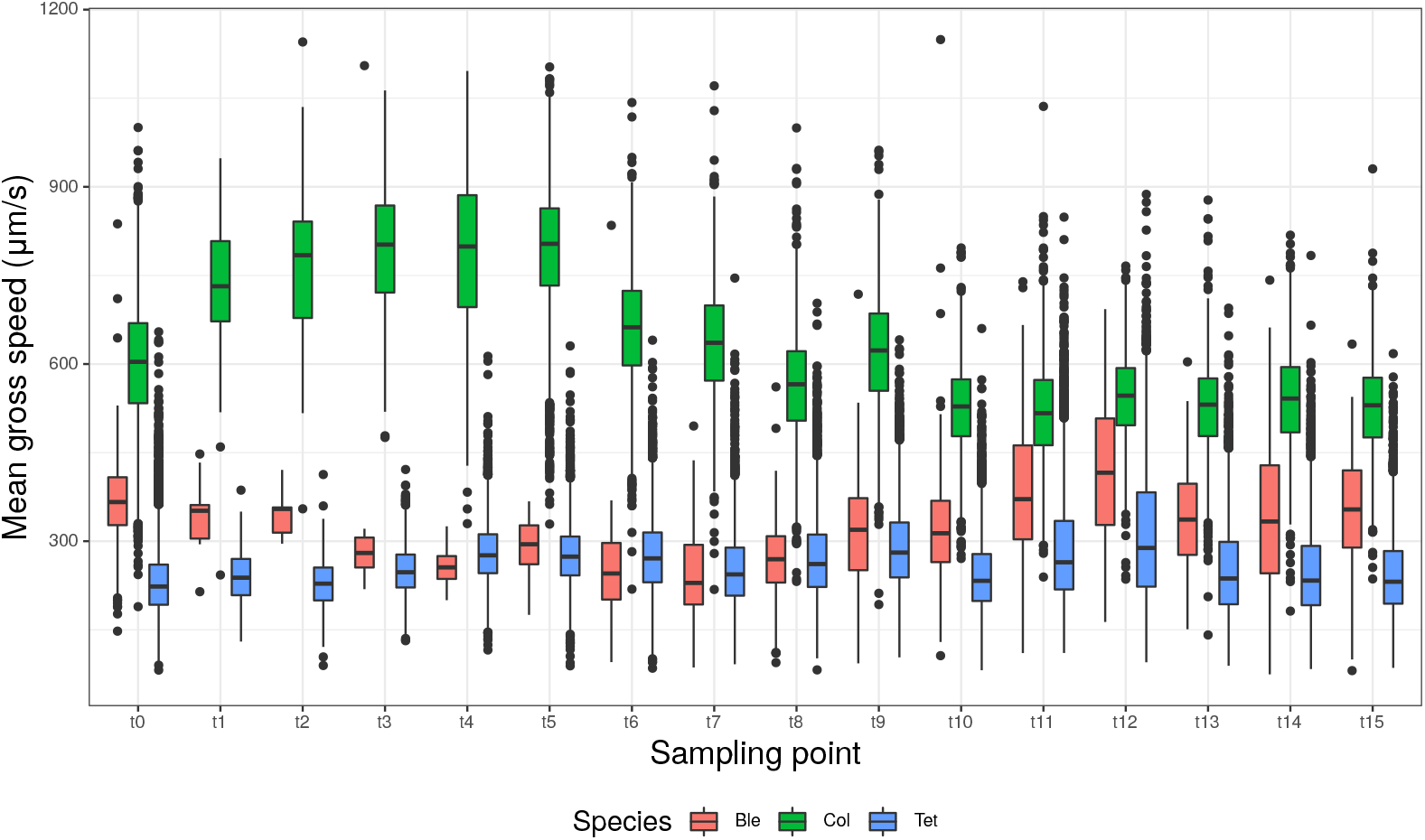
Gross speeds of *T. thermophila* (Tet), *Colpidium* sp. (Col) and *Blepharisma* sp. accross sampling points in single-patch mono-culutures.

**Figure S4.**
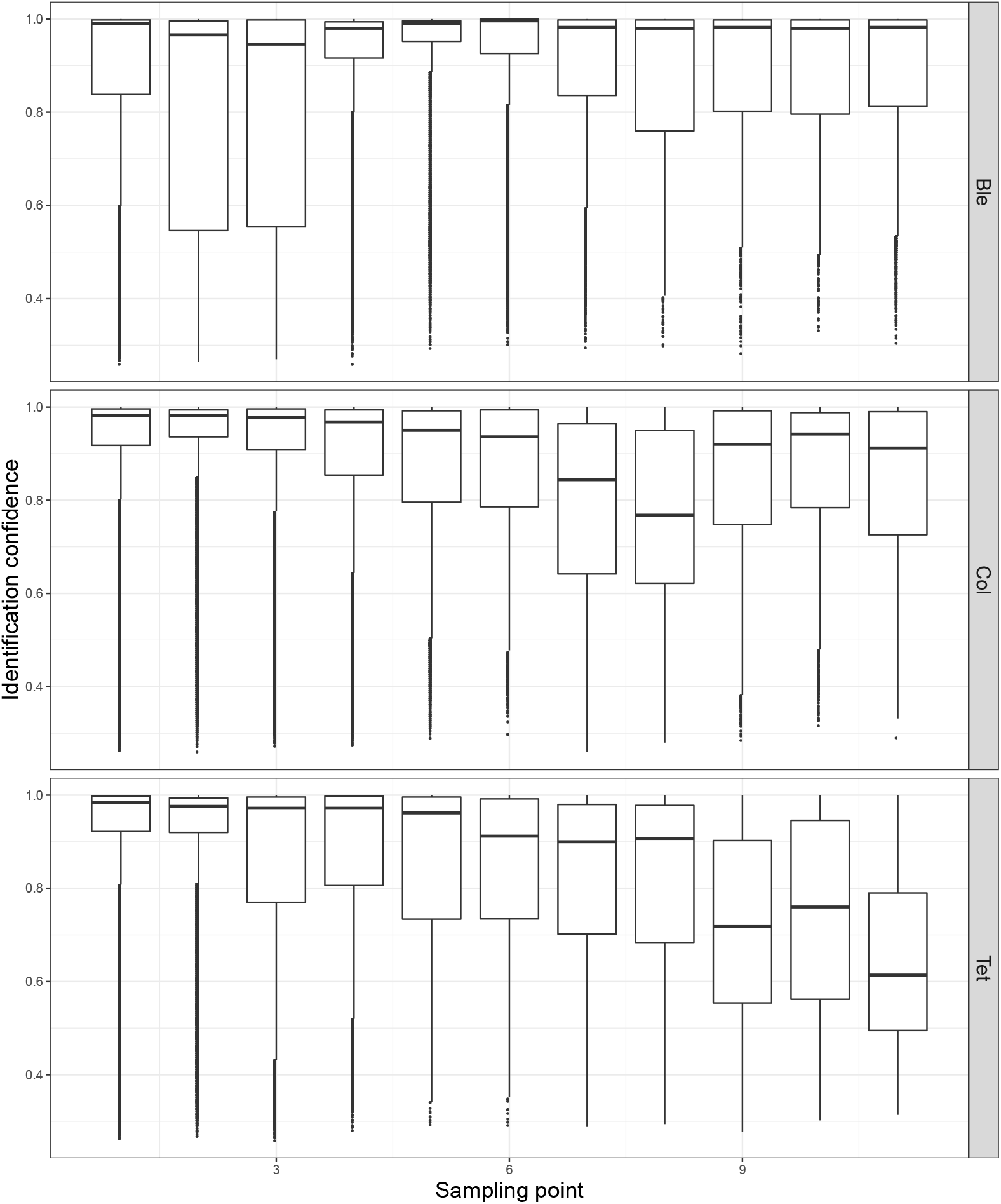
Identification confidence for individuals identified as *Blepharisma* sp. (Ble), *Colpidium* sp. and *T. Thermophila* at each sampling point of the experiment.

**Figure S5.**
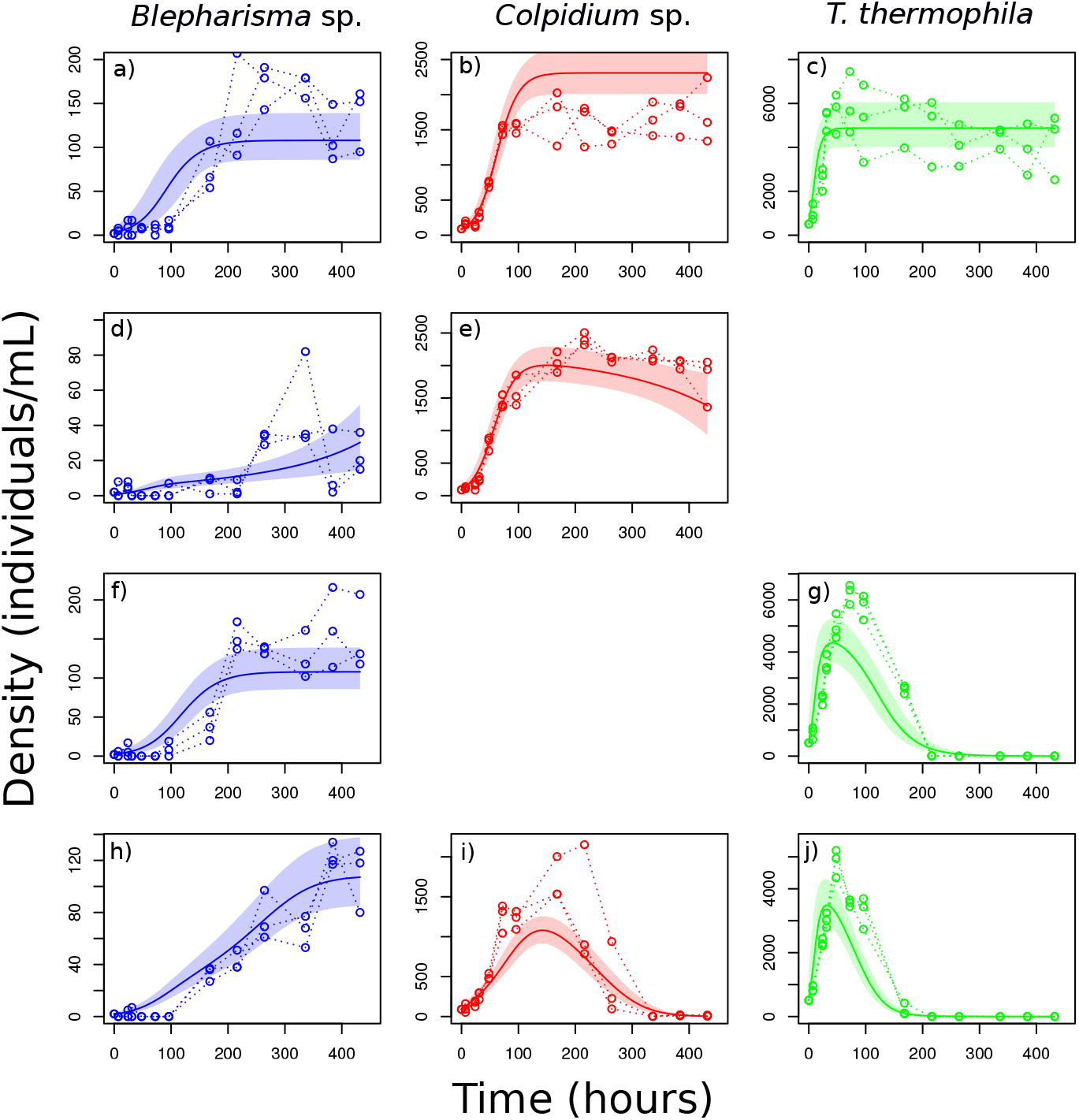
Fit of a competitive Lotka-Volterra model to experimental time series data obtained in single patch cultures of *Blepharisma* sp. (blue), *Colpidium* sp. (red) and *T. thermophila* (green). The curves and shaded areas show the posterior model predictions (median and 95% CI), the points and dashed lines show the experimental densities. The first line (a, b, c) shows the monoculture of each species. The second and third lines (d, e, f, g) show co-cultures of *Blepharisma* sp. with *Colpidium* sp. (d, e) and *Blepharisma* sp. with *T. thermophila* (f, g). The fourth line (h, i, j) shows the co-culture of all three species together.

**Figure S6.**
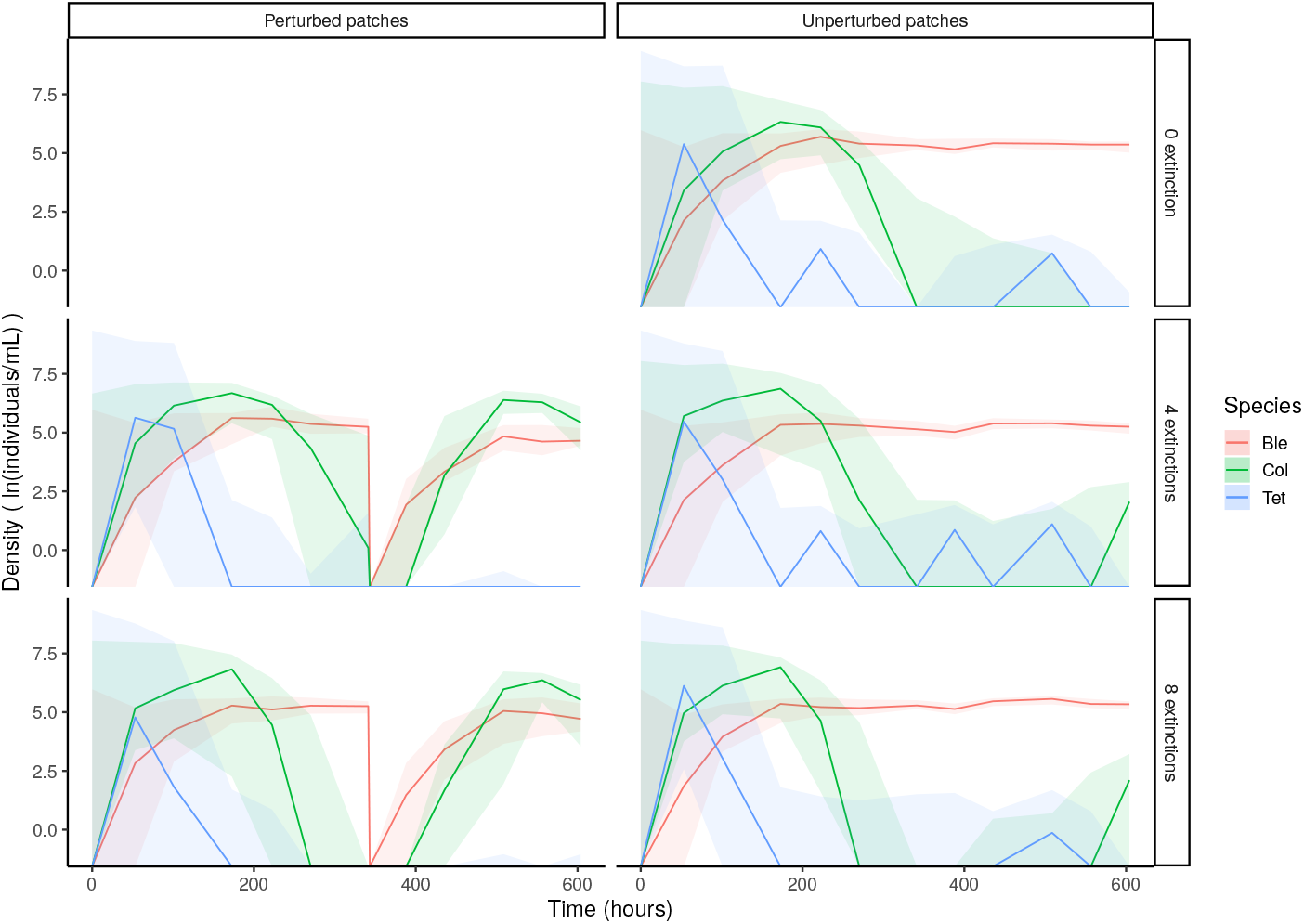
Median (solid line) and quantiles (colored areas) of species densities during the experiments (*Blepharisma* sp. in red, *Colpidium* sp. in green and *Tetrahymena termophila* in blue). The left column shows perturbed patches, in which *Blepharisma* sp. and *Colpidium* sp. had similar biomass during the recolonization process resulting in a high local diversity. The right column shows unperturbed patches from control landscapes (top) and from landscapes with extinctions (middle and bottom), in which *Blepharisma* sp. quickly became dominant, resulting in a low local diversity. Note that the scale of density is logarithmic.

**Figure S7.**
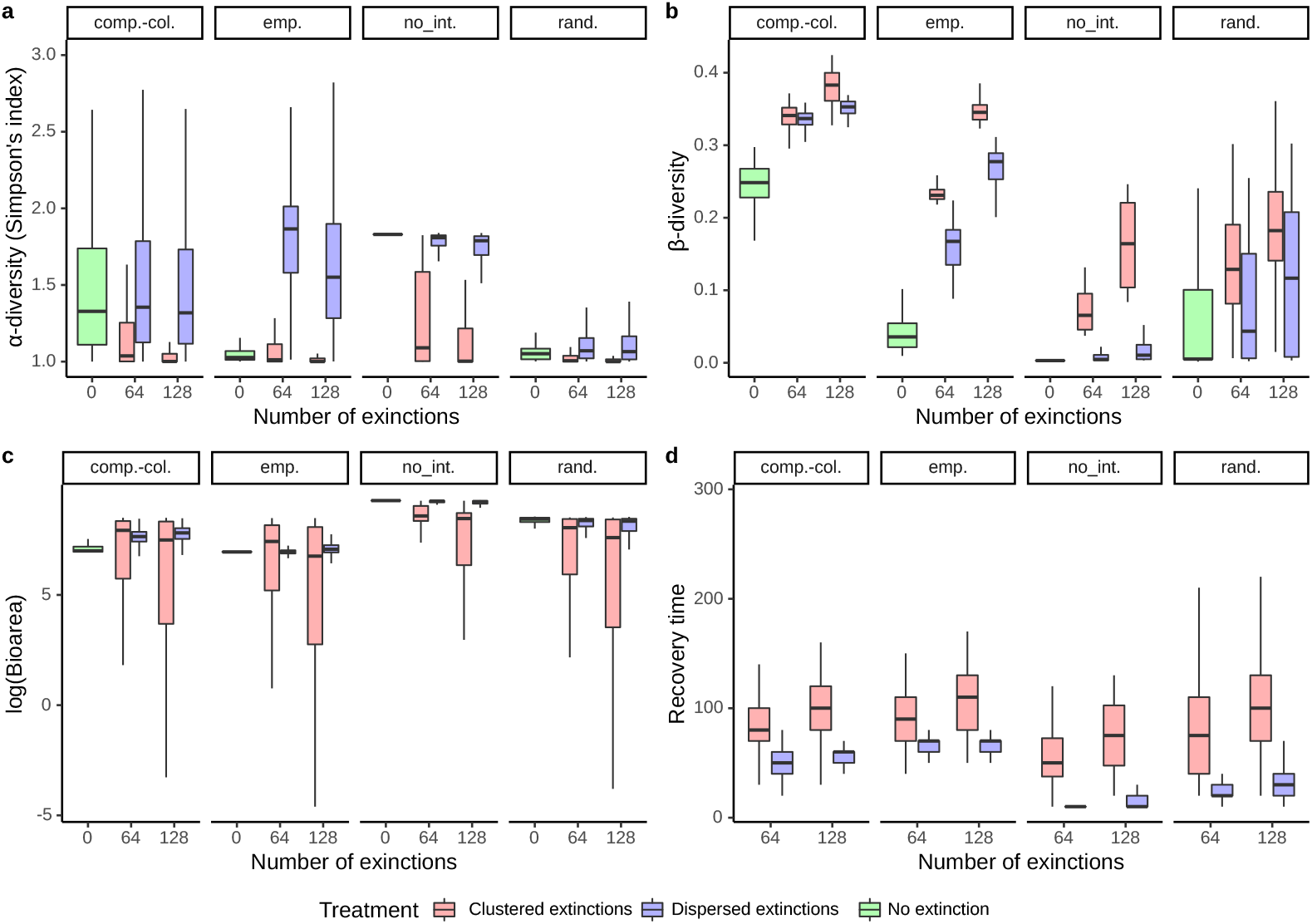
Sensitivity analysis: large landscape (16*16 patches). Observed response variables in numerical simulations of the metacommunity model displaying different metrics after the extinction events. (a) *α*-diversity (measured as Simpson’s index) in perturbed patches, (b) *β*-diversity in landscapes with extinction, (c) biomass in perturbed patches and (d) biomass recovery time in perturbed patches. The top labels denote the scenarios of species interactions: “emp.” for “empirical interactions”, “comp.-col.” for “competition-colonization trade-off”, “rand.” for “randomized interactions” and “no int.” for “no interspecific interactions”. Results are qualitatively similar to what was found in smaller landscapes – see Fig. 2 for comparison.

**Figure S8.**
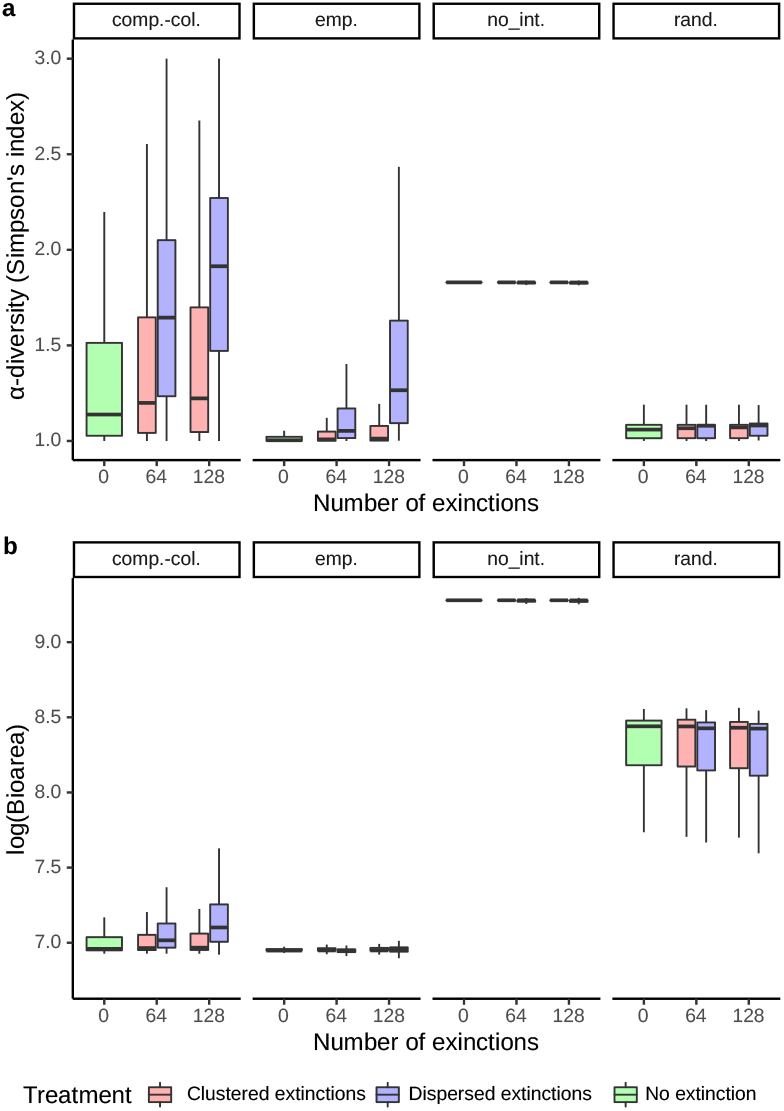
Sensitivity analysis: large landscape (16*16 patches). Observed response variables in numerical simulations of the metacommunity model showing Simpson’s index (a) and biomass (b) in unperturbed patches adjacent to at least one perturbed patches (blue, red) and in control landscapes (green) after extinction events. The top labels denote the scenarios of species interactions: “emp.” for “empirical interactions”, “comp.-col.” for “competition-colonization trade-off”, “rand.” for “randomized interactions” and “no int.” for “no interspecific interactions”. Results are qualitatively similar to what was found in smaller landscapes – see Fig. 4 for comparison.

**Figure S9.**
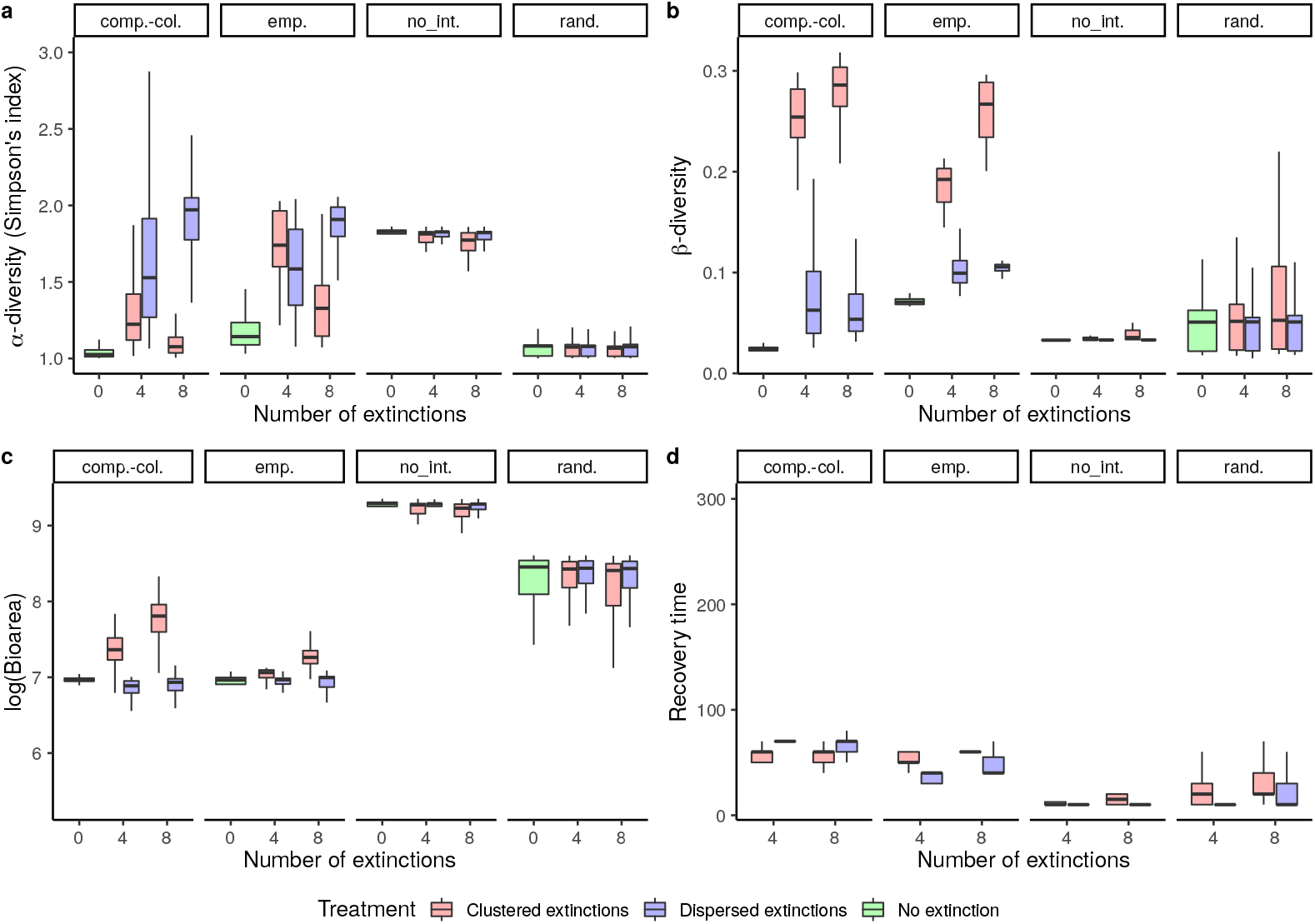
Sensitivity analysis: dispersal (5 times stronger). Observed response variables in numerical simulations of the metacommunity model displaying different metrics after the extinction events. (a) *α*-diversity (measured as Simpson’s index) in perturbed patches, (b) *β*-diversity in landscapes with extinction, (c) biomass in perturbed patches and (d) biomass recovery time in perturbed patches. The top labels denote the scenarios of species interactions: “emp.” for “empirical interactions”, “comp.-col.” for “competition-colonization trade-off”, “rand.” for “randomized interactions” and “no int.” for “no interspecific interactions”.

**Figure S10.**
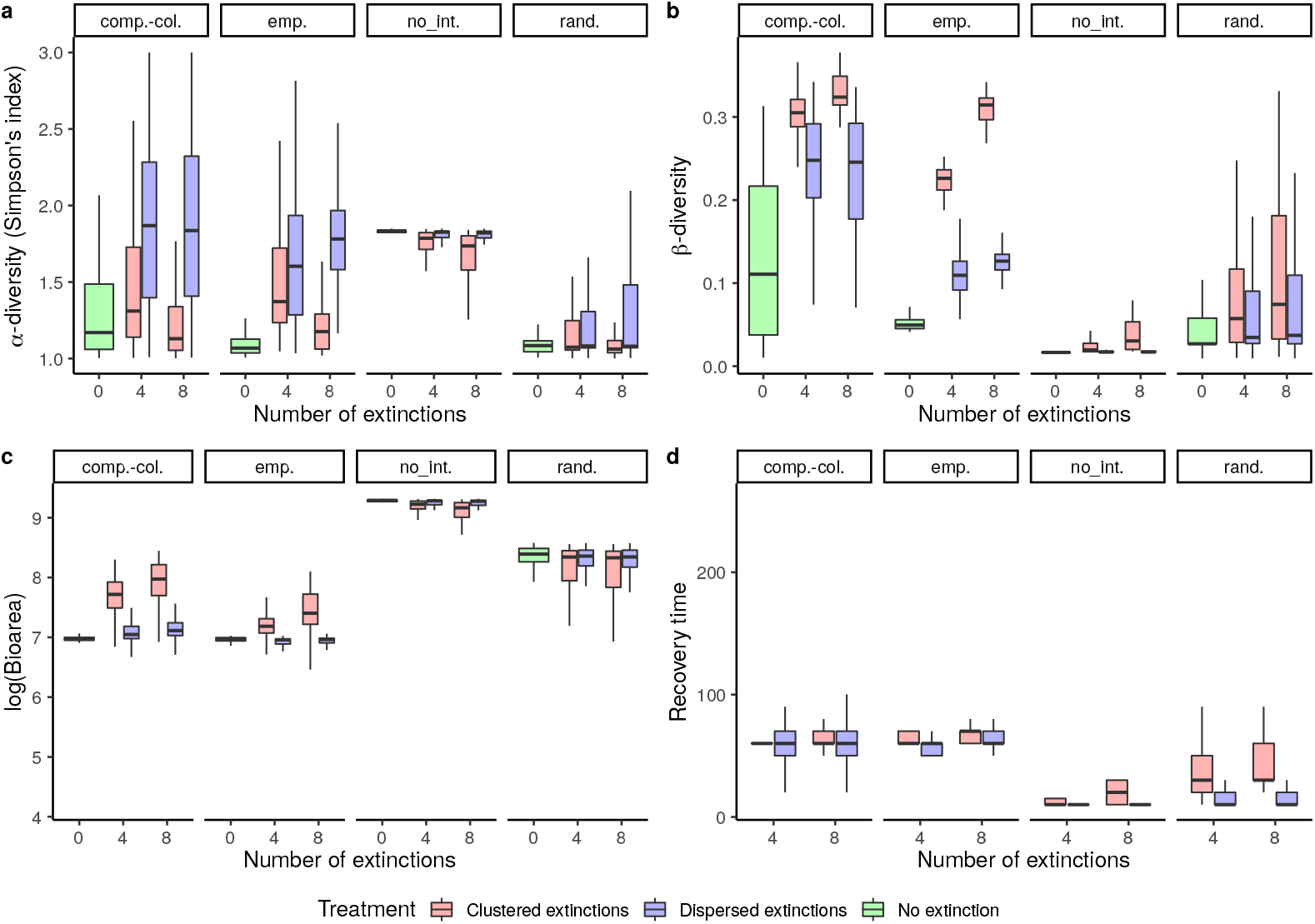
Sensitivity analysis: dispersal (2 times stronger). Observed response variables in numerical simulations of the metacommunity model displaying different metrics after the extinction events. (a) *α*-diversity (measured as Simpson’s index) in perturbed patches, (b) *β*-diversity in landscapes with extinction, (c) biomass in perturbed patches and (d) biomass recovery time in perturbed patches. The top labels denote the scenarios of species interactions: “emp.” for “empirical interactions”, “comp.-col.” for “competition-colonization trade-off”, “rand.” for “randomized interactions” and “no int.” for “no interspecific interactions”.

**Figure S11.**
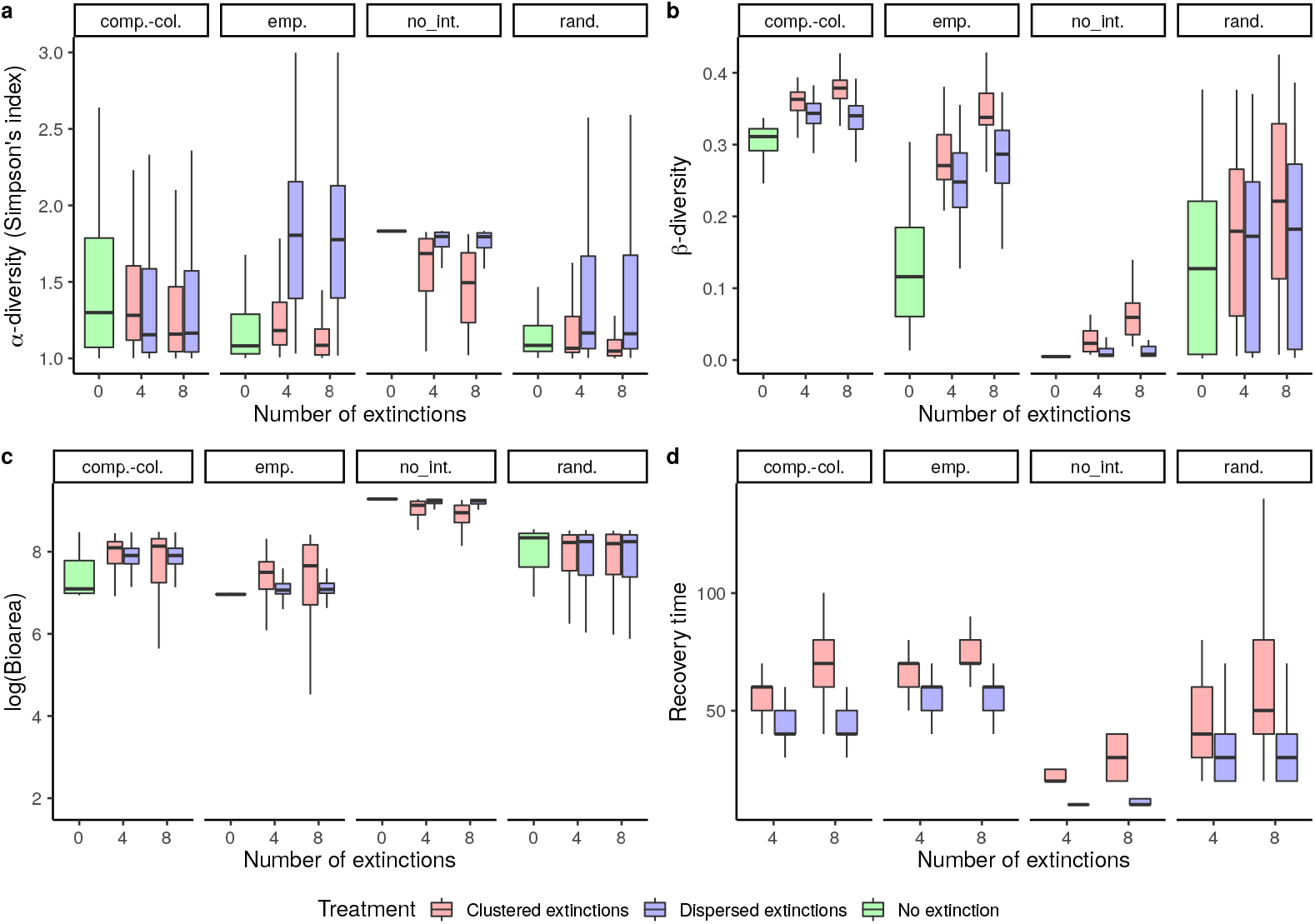
Sensitivity analysis: dispersal (2 times weaker). Observed response variables in numerical simulations of the metacommunity model displaying different metrics after the extinction events. (a) *α*-diversity (measured as Simpson’s index) in perturbed patches, (b) *β*-diversity in landscapes with extinction, (c) biomass in perturbed patches and (d) biomass recovery time in perturbed patches. The top labels denote the scenarios of species interactions: “emp.” for “empirical interactions”, “comp.-col.” for “competition-colonization trade-off”, “rand.” for “randomized interactions” and “no int.” for “no interspecific interactions”.

**Figure S12.**
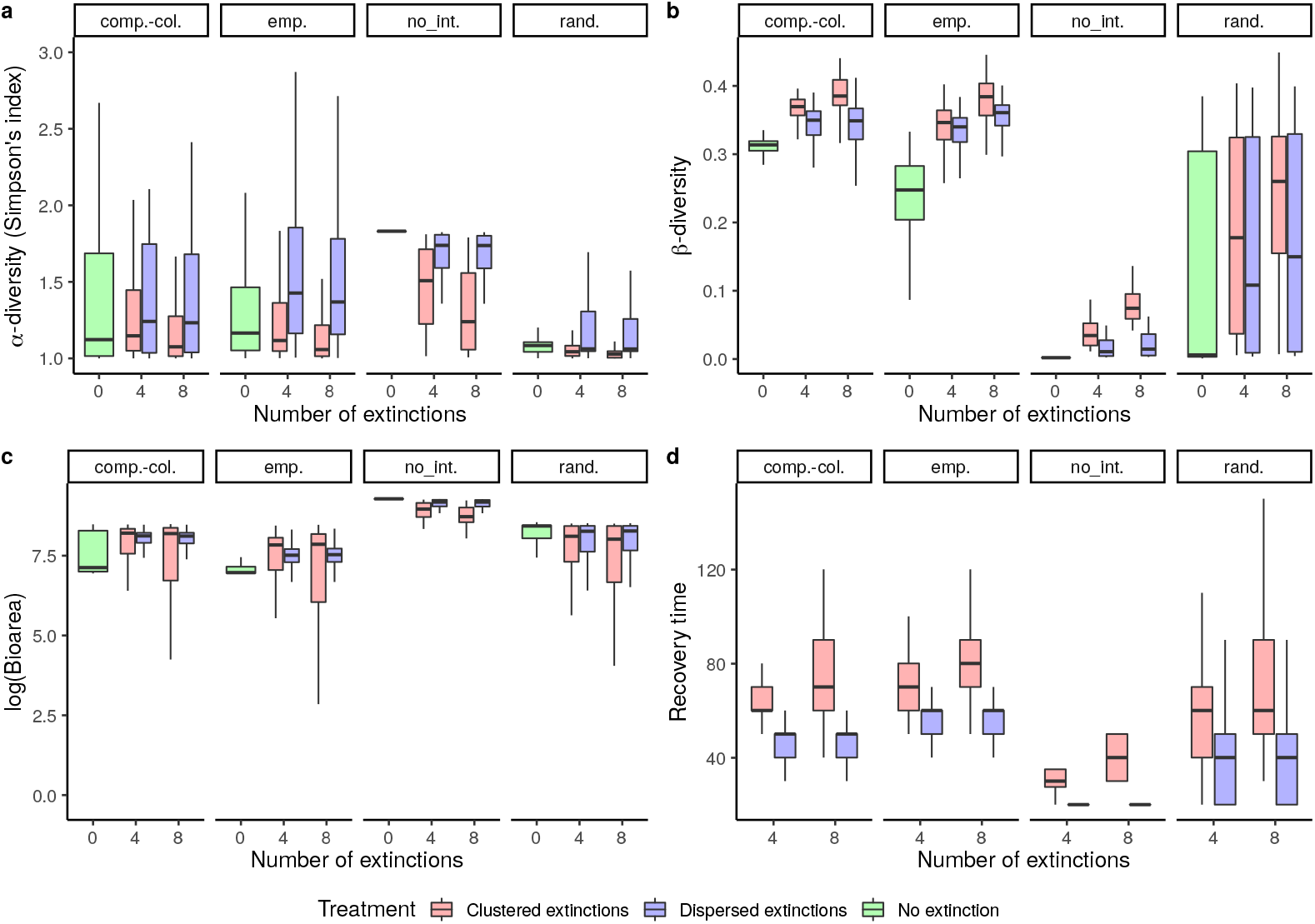
Sensitivity analysis: dispersal (5 times weaker). Observed response variables in numerical simulations of the metacommunity model displaying different metrics after the extinction events. (a) *α*-diversity (measured as Simpson’s index) in perturbed patches, (b) *β*-diversity in landscapes with extinction, (c) biomass in perturbed patches and (d) biomass recovery time in perturbed patches. The top labels denote the scenarios of species interactions: “emp.” for “empirical interactions”, “comp.-col.” for “competition-colonization trade-off”, “rand.” for “randomized interactions” and “no int.” for “no interspecific interactions”.

**Figure S13.**
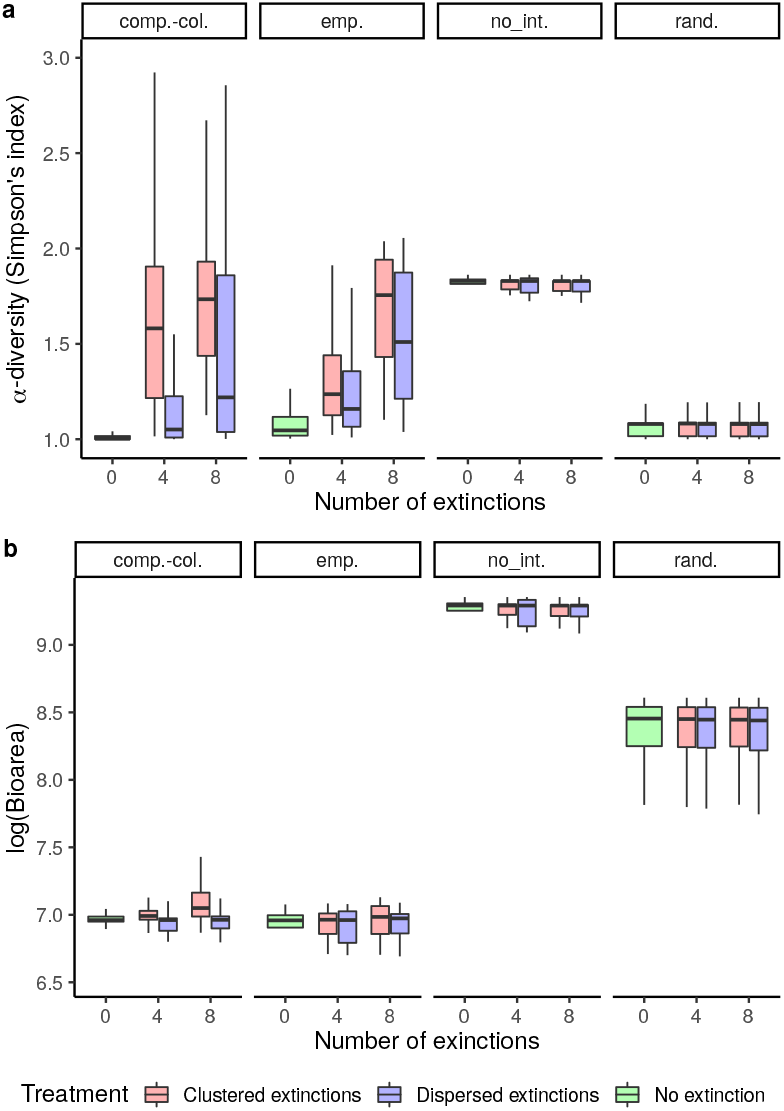
Sensitivity analysis: dispersal (5 times stronger). Observed response variables in numerical simulations of the metacommunity model showing Simpson’s index (a) and biomass (b) in unperturbed patches adjacent to at least one perturbed patches (blue, red) and in control landscapes (green) after extinction events. The top labels denote the scenarios of species interactions: “emp.” for “empirical interactions”, “comp.-col.” for “competition-colonization trade-off”, “rand.” for “randomized interactions” and “no int.” for “no interspecific interactions”.

**Figure S14.**
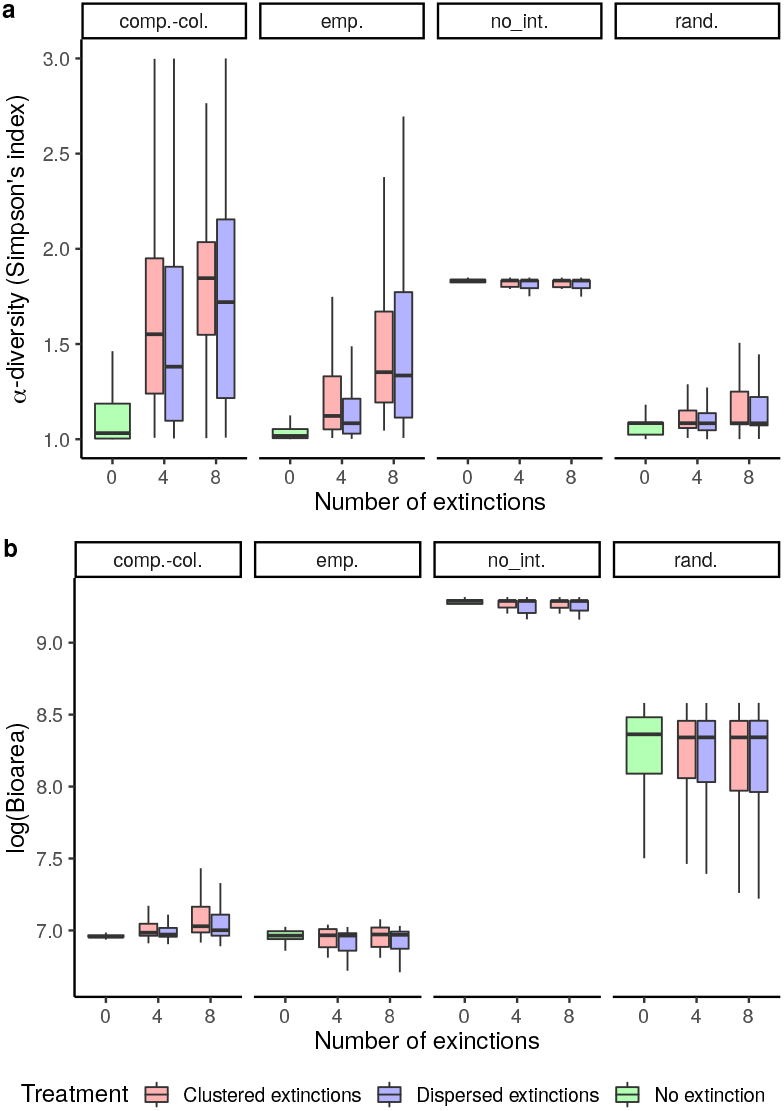
Sensitivity analysis: dispersal (2 times stronger). Observed response variables in numerical simulations of the metacommunity model showing Simpson’s index (a) and biomass (b) in unperturbed patches adjacent to at least one perturbed patches (blue, red) and in control landscapes (green) after extinction events. The top labels denote the scenarios of species interactions: “emp.” for “empirical interactions”, “comp.-col.” for “competition-colonization trade-off”, “rand.” for “randomized interactions” and “no int.” for “no interspecific interactions”.

**Figure S15.**
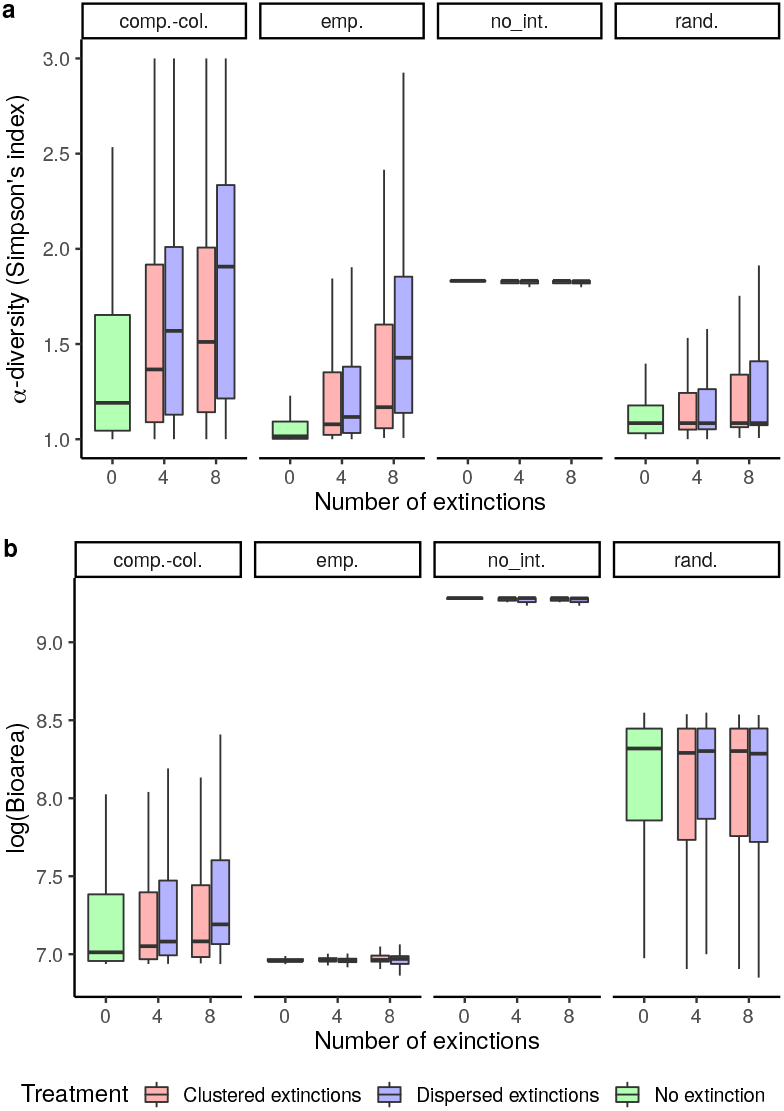
Sensitivity analysis: dispersal (2 times weaker). Observed response variables in numerical simulations of the metacommunity model showing Simpson’s index (a) and biomass (b) in unperturbed patches adjacent to at least one perturbed patches (blue, red) and in control landscapes (green) after extinction events. The top labels denote the scenarios of species interactions: “emp.” for “empirical interactions”, “comp.-col.” for “competition-colonization trade-off”, “rand.” for “randomized interactions” and “no int.” for “no interspecific interactions”.

**Figure S16.**
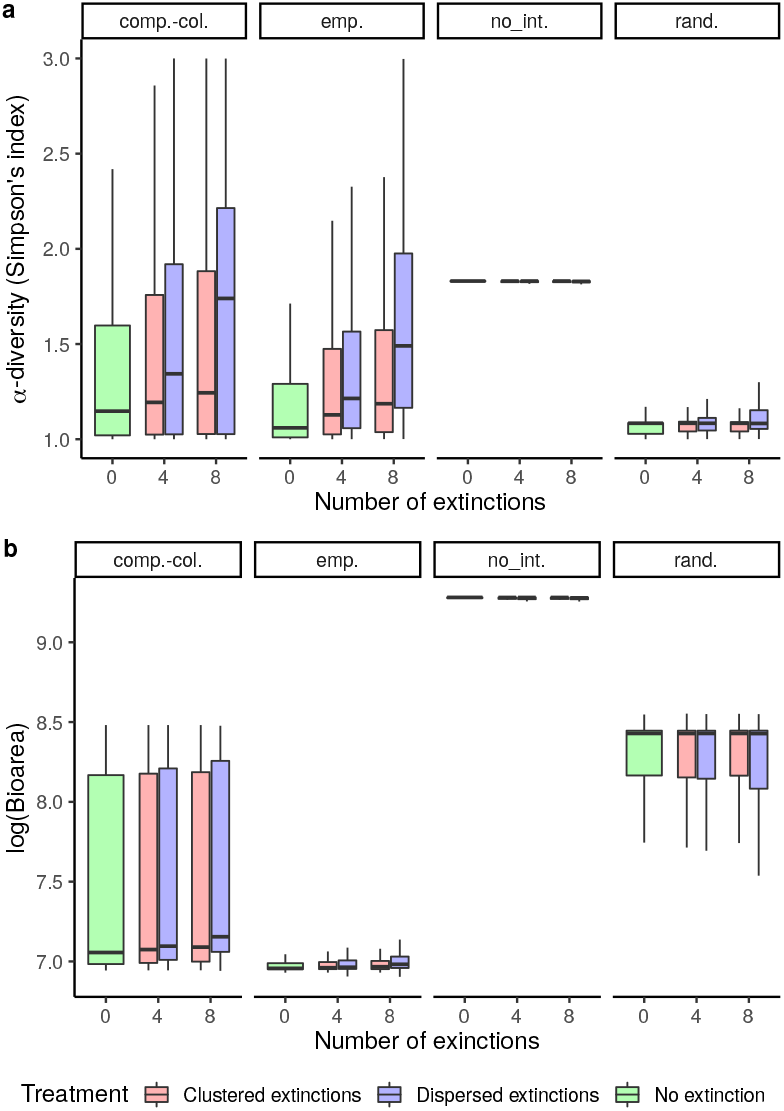
Sensitivity analysis: dispersal (5 times weaker). Observed response variables in numerical simulations of the metacommunity model showing Simpson’s index (a) and biomass (b) in unperturbed patches adjacent to at least one perturbed patches (blue, red) and in control landscapes (green) after extinction events. The top labels denote the scenarios of species interactions: “emp.” for “empirical interactions”, “comp.-col.” for “competition-colonization trade-off”, “rand.” for “randomized interactions” and “no int.” for “no interspecific interactions”.

**Figure S17.**
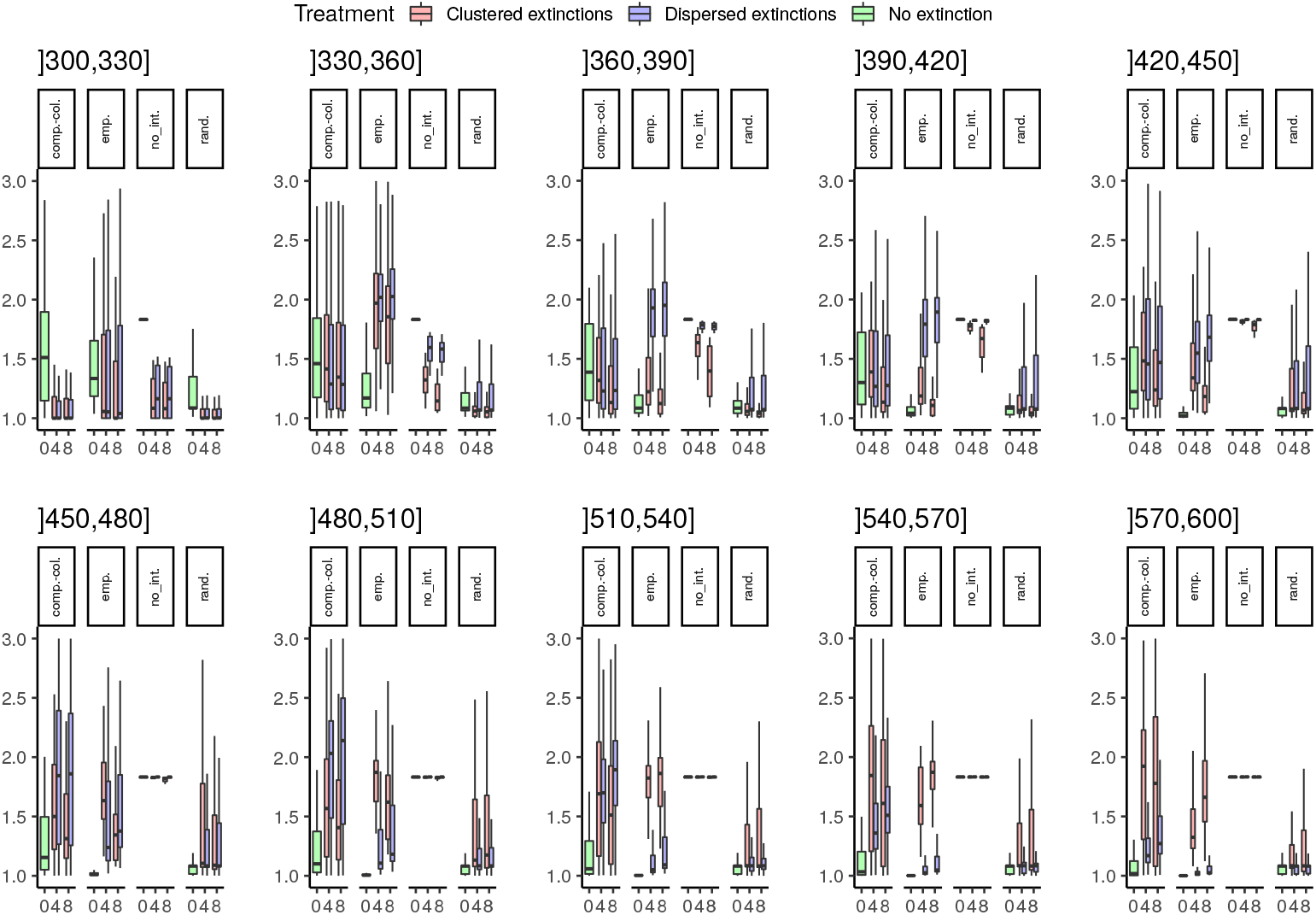
*α*-diversity in perturbed patches in numerical simulations of the metacommunity model over moving time windows from the extinction time (300 time units) to the end of the simulation (600 time units). The bottom labels denote the amount of extinctions (0, 4, 8). The top labels denote the scenarios of species interactions: “emp.” for “empirical interactions", “comp.-col.” for “competition/colonization trade-off", “rand.” for “randomized interactions” and “no_int.” for “no interspecific interactions”.

**Figure S18.**
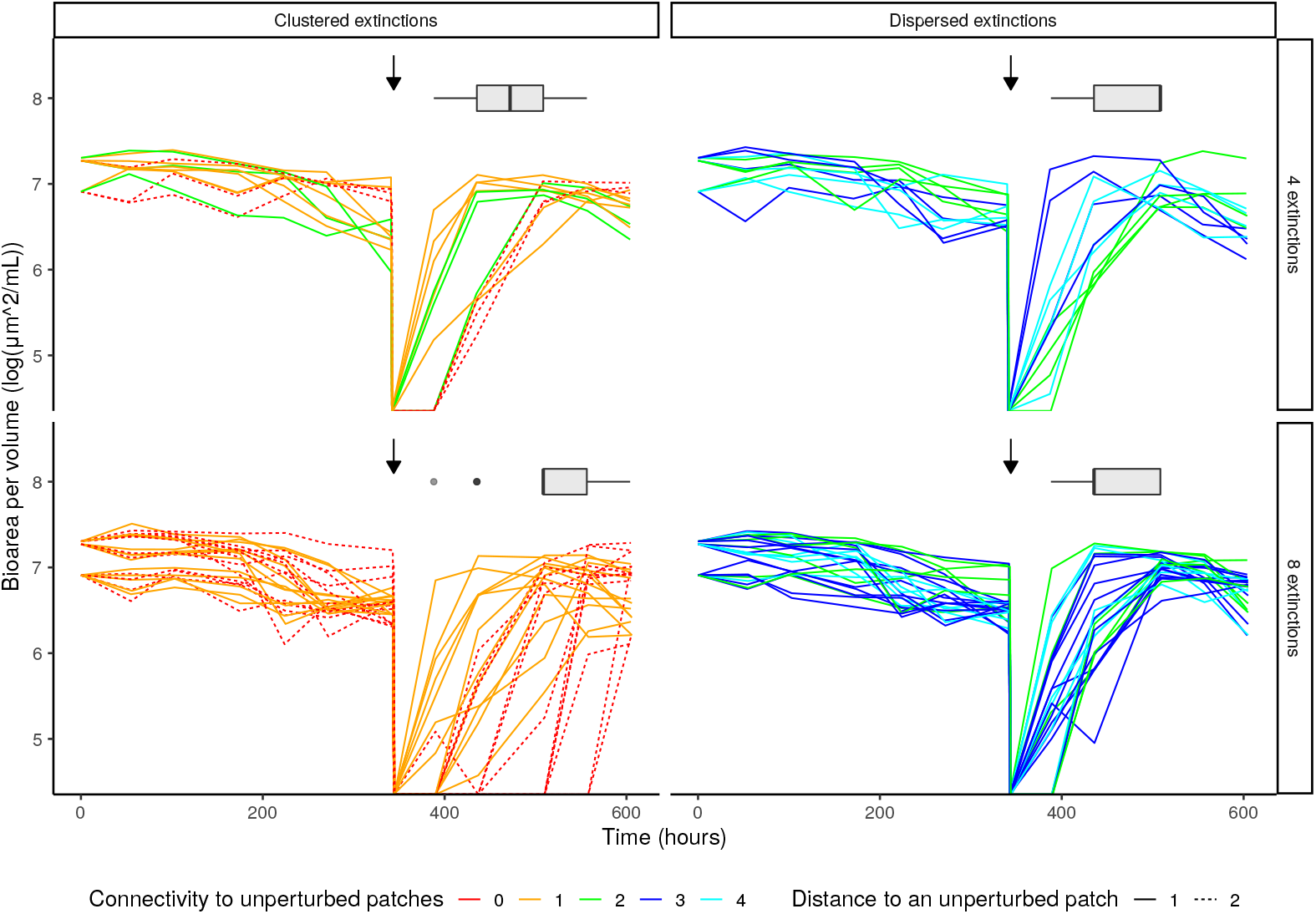
Bioarea over time in perturbed patches. Each panel represents a treatment (columns: spatial autocorrelation of extinctions; lines: amount of extinctions). The vertical arrows show the time at which extinctions happened. The boxes represent the distribution of recovery times (time needed to reach a bioarea per volume higher that the 2.5% quantile of pre-extinction bioarea in a given patch) in each treatment. Dashed lines indicate that the patch is not directly in contact to an unperturbed patch (distance of 2 connections). The colors indicate the number of adjacent unperturbed patches (cyan : 4, blue : 3, green : 2, orange : 1 and red : 0).

**Figure S19.**
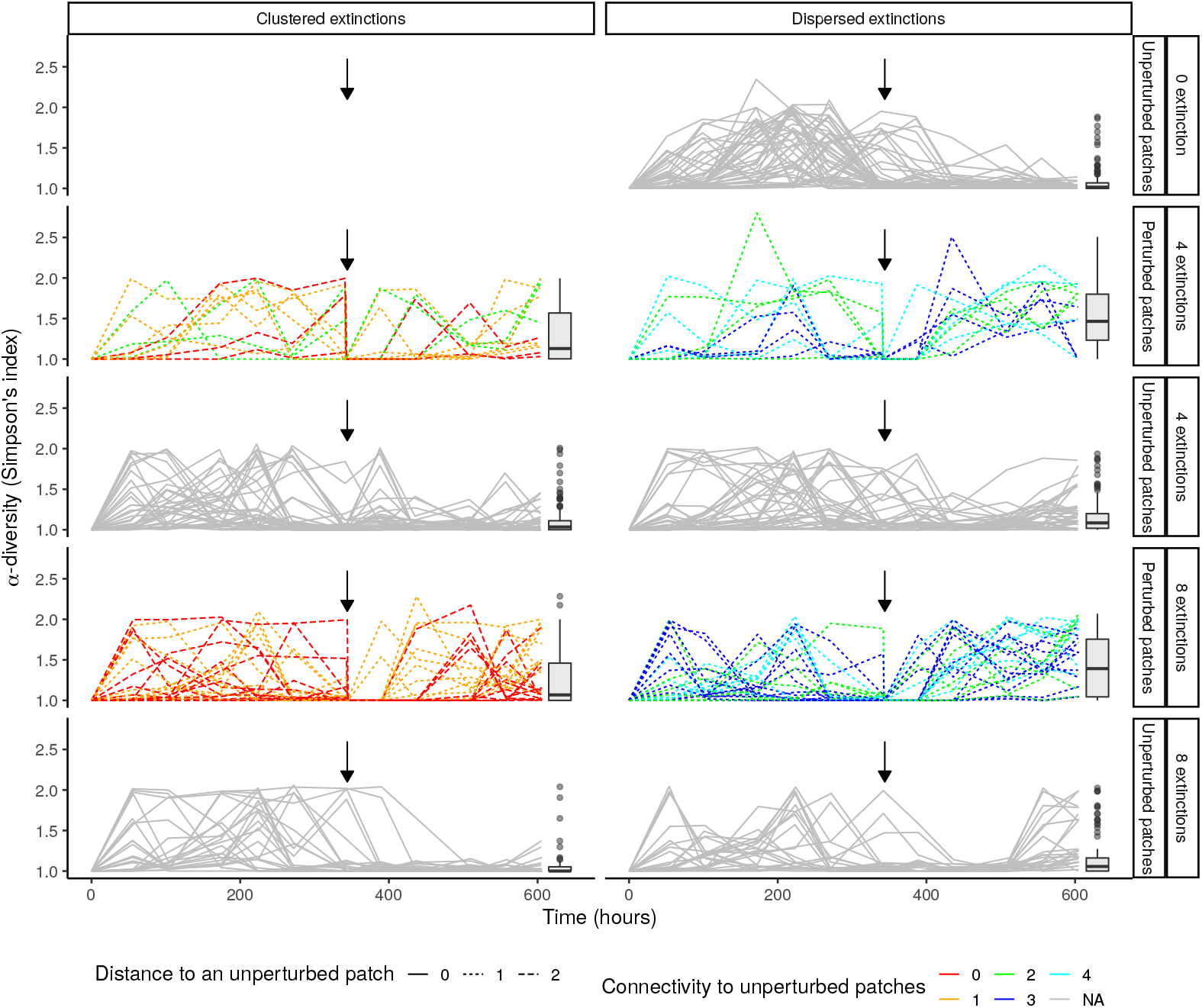
*α*-diversity over time in all patches. Each panel represents a treatment (columns: spatial autocorrelation of extinctions; lines: amount of extinctions). The vertical arrows show the time at which extinctions happened. The boxes represent the distribution of *α*-diversity after the extinctions.

**Figure S20.**
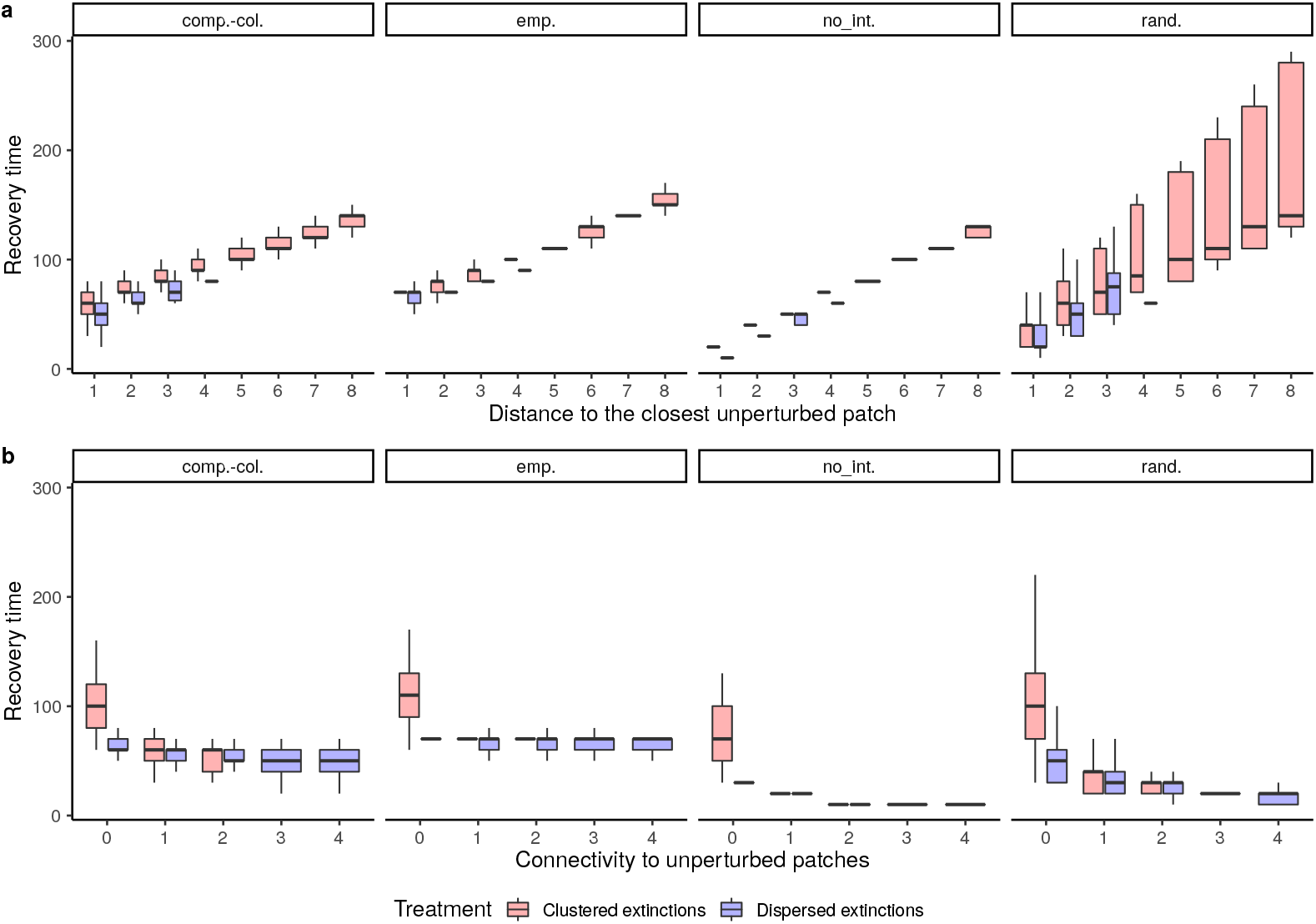
Recovery time (time needed to reach a bioarea per volume higher that the 2.5% quantile of pre-extinction bioarea in a given patch) of extinct patches as a function of (a) the distance to the closest unperturbed patches and (b) the connectivity to unperturbed patches (*i.e.*, the number of adjacent unperturbed patches), in simulations of large (16 × 16) landscapes. The top labels denote the scenarios of species interactions: “emp.” for “empirical interactions”, “comp.-col.” for “competition-colonization trade-off”, “rand.” for “randomized interactions” and “no int.” for “no interspecific interactions”.

**Figure S21.**
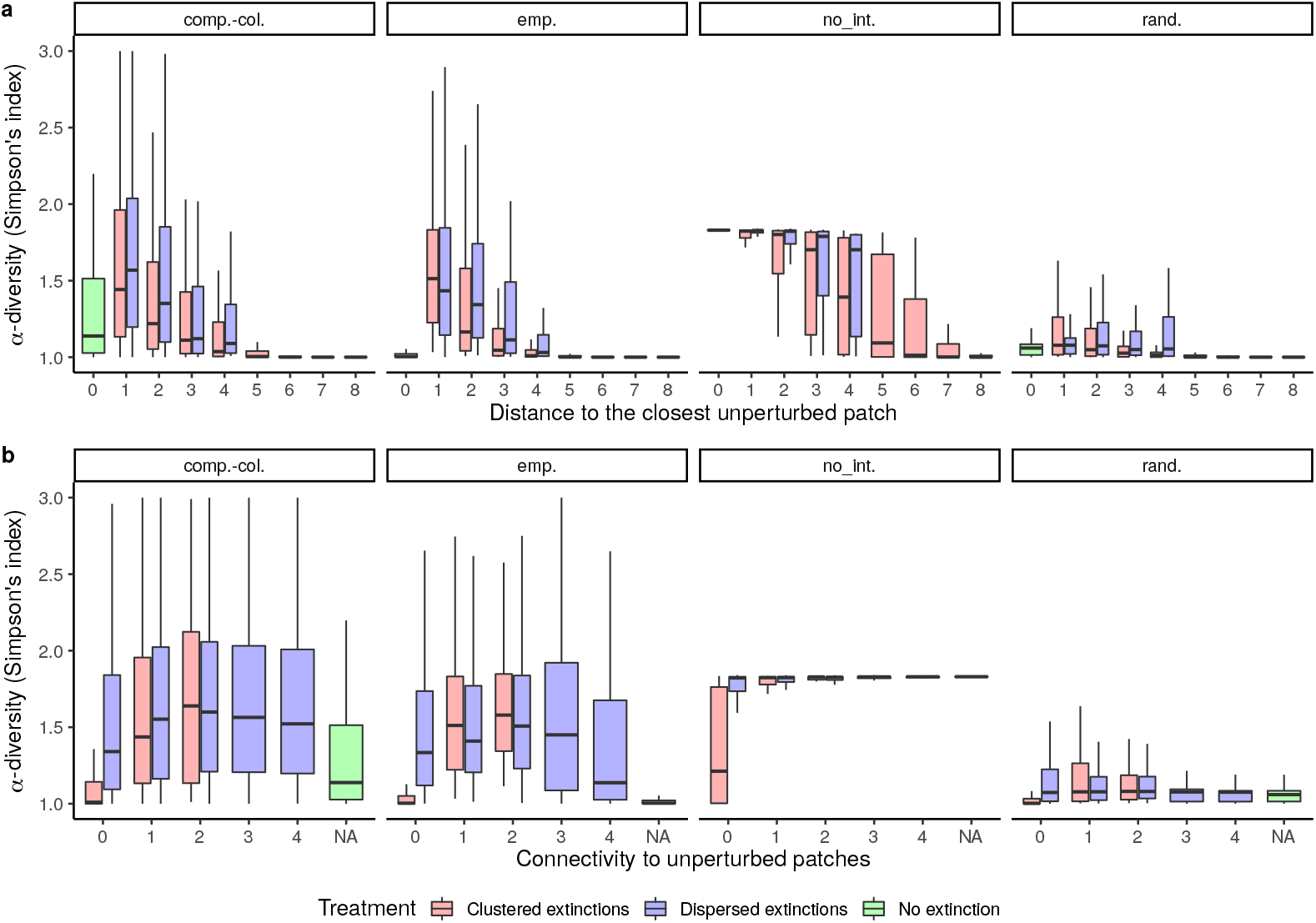
*α*-diversity of extinct patches (red, blue) during the recolonization process as a function of (a) the distance to the closest unperturbed patches and (b) the connectivity to unperturbed patches (*i.e.*, the number of adjacent unperturbed patches), in simulations of large (16×16) landscapes. The colors denote the spatial treatment (red: clustered extinctions, blue: dispersed extinctions). Patches from control landscapes (green: no extinction) where added as a reference and assigned a distance of 0 and a “NA” for connectivity. The top labels denote the scenarios of species interactions: “emp.” for “empirical interactions”, “comp.-col.” for “competition-colonization trade-off”, “rand.” for “randomized interactions” and “no int.” for “no interspecific interactions”.

**Figure S22.**
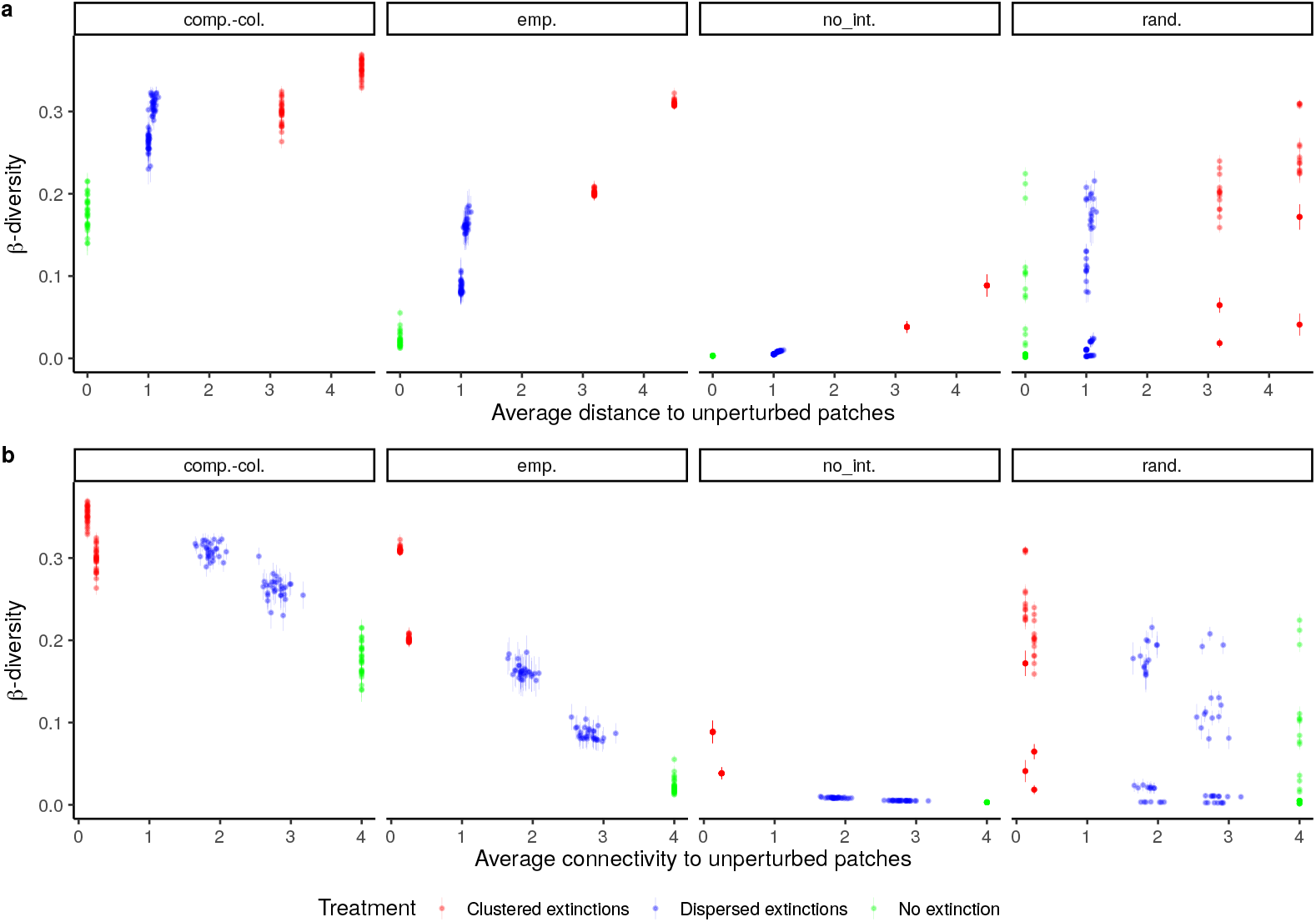
*β*-diversity of all landcapes during the recolonization process as a function of (a) the average distance to the closest unperturbed patch and (b) the average connectivity to unperturbed patches (*i.e.*, the number of adjacent unperturbed patches), in simulations of large (16×16) landscapes. The colors denote the spatial treatment (red: clustered extinctions, blue: dispersed extinctions, green: no extinction). The top labels denote the scenarios of species interactions: “emp.” for “empirical interactions”, “comp.-col.” for “competition-colonization trade-off”, “rand.” for “randomized interactions” and “no int.” for “no interspecific interactions”.

### Supplementary Tables

**Table S1.** Priors used to fit a competitive Lotka-Volterra model on experimental time series. We used the same growth rates (*r_i_*, one per species) and competition strengths (one intraspecific term (*α_i,i_*) per species and 6 interspecific terms (*α_i,j_*_; *i*≠*j*_)) over all replicates. We fitted unique initial densities (*N*_0_) on each species in each replicate.

| Parameters | Meaning | prior |
| --- | --- | --- |
| $r_i$ | Growth rates | lognormal(-2, 1) |
| $\alpha_{i,j}$ | Competition strengths | gamma(2, 1) |
| $N_0$ | Initial densities | *Blepharisma* sp.: normal(0,10) |
|  |  | *Colpidium* sp.: normal(0,100) |
|  |  | *T. thermophila*: normal(0,1000) |

**Table S2.** Average properties of perturbed patches across treatments: connectivity to unperturbed patches (*i.e.*, the number or unperturbed adjacent patches) and distance to the closest unperturbed patch.

| Treatment: | 4 clustered | 4 dispersed | 8 clustered | 8 dispersed |
| --- | --- | --- | --- | --- |
| Distance to an unperturbed patch | 1.25 | 1 | 1.5 | 1 |
| Connectivity to unperturbed patches | 1 | 3 | 0.5 | 3 |

**Table S3.** Tables of model comparison for local effects in perturbed patches (*α*-diversity, bioarea and recovery time) and *β*-diversity. For each variable, we compared all mixed models between the full model (Spatial autocorrelation * Amount of extinctions) and the intercept using AICc. Models not displayed – for *β*-diversity (b) – had a negligible weight.

| Model | $\Delta AICc$ | Weight |
| --- | --- | --- |
| Spatial autocorrelation | 0.00 | 0.542 |
| Spatial autocorrelation + Amount of extinctions | 1.15 | 0.302 |
| Spatial autocorrelation * Amount of extinctions | 3.23 | 0.108 |
| Intercept | 5.71 | 0.031 |
| Amount of extinctions | 7.16 | 0.015 |

**(b)** Beta diversity model comparison.
| Model | $\Delta AICc$ | Weight |
| --- | --- | --- |
| Spatial autocorrelation * Amount of extinctions | 0.00 | 1 |
| Spatial autocorrelation + Amount of extinctions | 20.44 | 0.00 |

**(c)** Bioarea model comparison.
| Model | $\Delta AICc$ | Weight |
| --- | --- | --- |
| Spatial autocorrelation | 0.00 | 0.362 |
| Intercept | 0.62 | 0.265 |
| Spatial autocorrelation + Amount of extinctions | 1.70 | 0.155 |
| Amount of extinctions | 2.32 | 0.113 |
| Spatial autocorrelation * Amount of extinctions | 2.48 | 0.105 |

**(d)** Recovery time model comparison.
| Model | $\Delta AICc$ | Weight |
| --- | --- | --- |
| Spatial autocorrelation | 0.00 | 0.271 |
| Spatial autocorrelation * Amount of extinctions | 0.31 | 0.232 |
| Intercept | 0.50 | 0.211 |
| Spatial autocorrelation + Amount of extinctions | 0.93 | 0.170 |
| Amount of extinctions | 1.69 | 0.117 |

**Table S4.**
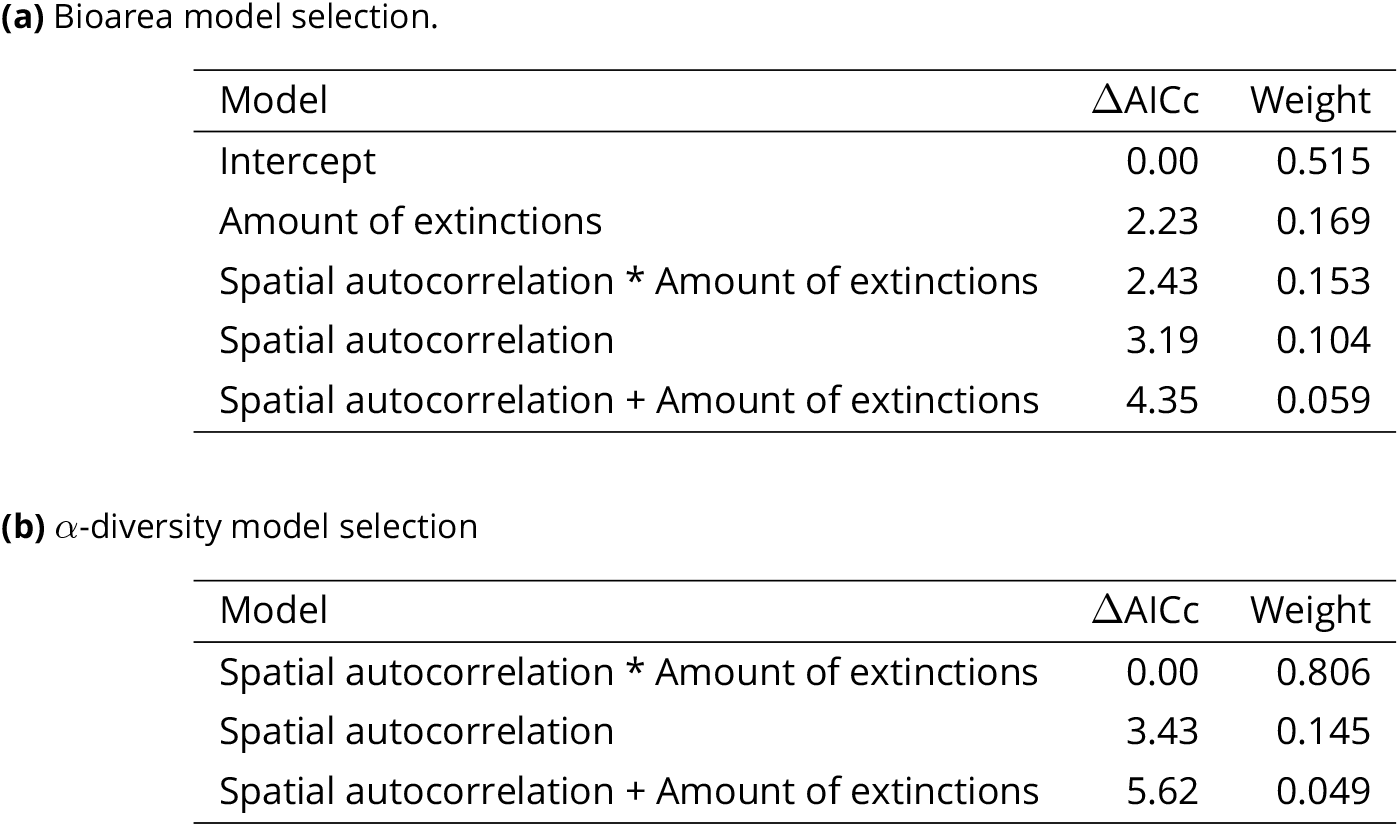
Tables of model comparison for bioarea and *α*−diversity in unperturbed patches adjacent to at least one perturbed patch. For both variables, we compared all mixed models between the full model (Spatial autocorrelation * Amount of extinctions) and the intercept using AICc. Models not displayed – for *α*-diversity (b) – had a negligible weight.

## Notes

https://doi.org/10.5281/zenodo.4660016

### Competing Interest Statement

The authors have declared no competing interest.

### Summary of Updates

Version 4 of this preprint has been peer-reviewed and recommended by Peer Community In Ecology (https://doi.org/10.24072/pci.ecology.100084)

